# Mental Chronometry of Speaking in Dialogue: Semantic Interference Turns into Facilitation

**DOI:** 10.1101/2020.09.08.287458

**Authors:** Anna K. Kuhlen, Rasha Abdel Rahman

## Abstract

Numerous studies demonstrate that the production of words is delayed when speakers process in temporal proximity semantically related words. Yet the experimental settings underlying this effect are different from those under which we typically speak. This study demonstrates that semantic interference disappears, and can even turn into facilitation, when semantically related words are embedded in a meaningful communicative exchange. Experiment 1 and 3 (each N=32 university students) implemented a picture-word interference task in a game played between two participants: one named the distractor word and, after a stimulus-onset-asynchrony of -150ms or -650ms, the other named a semantically related or unrelated target picture. Semantic interference reappeared with identical experimental parameters in a single-person picture-word interference setting (Experiment 2, N=32). We conclude that the inhibitory context effects leading to semantic interference in single-subject settings are attenuated whereas facilitatory effects are enhanced in communicative settings.

**Highlights:**

- In isolated speakers processing semantically related words interferes with speaking
- Yet we typically speak to communicate with others
- This study transfers picture-word interference to a communicative setting
- Semantically related words produced by a partner do not interfere with speaking

---

Over the last decades a multitude of experimental studies have demonstrated that semantic context, typically presented in the form of words embedded in simple picture naming tasks, can interfere with language production. The present study suggests that semantic interference may be mostly present in settings in which single words are produced devoid of a communicative purpose. When semantic context is embedded in a communicative social interaction interference can turn into facilitation.

A central paradigm for studying semantic context effects is the picture-word interference (PWI) task. In this task speakers name target pictures while trying to ignore visually or auditorily presented distractor words. One very robust finding is semantic interference: pictures are named with a greater latency if they are named in close temporal proximity to a semantically related distractor word (e.g., Damian & Martin, 1999; Glaser & Düngelhoff, 1984; Schriefers et al., 1990). This effect has motivated competitive theories of lexical selection (e.g., Levelt et al., 1999; Roelofs, 2018; Schriefers et al., 1990; but see e.g., Finkbeiner & Caramazza, 2006; Mahon et al., 2007 for an alternative, non-competitive account). These competitive theories assume that, on the lexical level, distractors cause interference and delay target production since distractor and a semantically related target compete for selection. However, on a conceptual level, distractors may also function as a prime and facilitate the activation of the target by pre-activating the target concept and features that are shared between semantically related distractor and target. Evidence for such conceptual priming has been found in PWI studies when the distractor was presented relatively long before the target picture (e.g., Alario et al., 2000; Bloem et al., 2004; Glaser & Düngelhoff, 1984).

Some theories assuming lexical competition have therefore proposed that overall semantic context effects arise from a trade-off between interference due to competition at the lexical level and concurrent semantic facilitation due to priming at the conceptual level (Abdel Rahman & Melinger, 2019; 2009). According to this proposal whether overall facilitation or interference is observed depends on whether conceptual processing (priming) or lexical processing (competition) dominates. Here we propose that communicative settings may influence this trade-off by swaying the distractor effects in the PWI task towards meaning-dominant processing at the conceptual level.

The picture-word interference task has been very instrumental to insights into the architecture underlying word production. Yet, in day-to-day conversations interlocutors routinely produce words from the same semantic category. For example, at a restaurant, a party of guests might discuss the menu options (e.g., DuBois, John, & Englebretson, 2004). One may suggest ordering a classic chef salad, while the other suggests a pasta salad. Do these interlocutors experience interference from each other’s references to different types of dishes? Based on the substantial empirical support from picture-word interference studies, speakers should be experiencing interference when seeking lexical access to semantically related items in close temporal proximity. Indeed, in everyday conversation the time that elapses between the end of one speaker’s turn and the beginning of the next speaker’s turn is typically around 200ms (Stivers et al., 2009), a time range within which, in experimental settings, semantic interference can be observed. Yet, under the premise that dialogue is the primary site of language use (e.g., Clark, 1996; Levinson, 1983) a speaker’s cognitive system should be optimally equipped to speak in such settings. Experiencing semantic interference from an interlocutor’s previous turn would undermine fluent conversation.

The currently existing accounts of speech production are primarily based on observations made in experimental settings devoid of a social or communicative context. To date it is an open question how these accounts translate to communicative language use. Specifically, producing language within in a larger meaningful context, for instance within discourse or during a conversation about a specific topic, has not been integrated into theoretical models of language production, and we can therefore only speculate about how social-communicative situations may affect language production in typical semantic context paradigms. An architecture that includes largely parallel conceptual and lexical processing and a context-dependent trade-off between effects of conceptual priming and lexical interference provides the basis for such modulations (for a discussion, see Abdel Rahman & Melinger, 2019). Social context may modulate the trade-off towards stronger effects of meaning-based conceptual priming relative to lexical competition: When producing language as part of a communicative exchange, processing the meaning of the partner’s in relation to one’s own utterances is in the foreground. Thus, the conceptual relationship between interlocutors’ utterances is likely to be more relevant than in traditional laboratory settings with single participants. In the picture-word interference paradigm this could lead to stronger effects of distractor words on the conceptual level during speech planning. This could sway semantic context effects towards conceptual priming and thus facilitate word retrieval when the interlocutors’ utterances share a given semantic context.

In line with these assumptions, available empirical evidence suggests a high degree of flexibility during conceptual and lexical processing triggered by the larger semantic context. For instance, while associative relations in the PWI task induce faster naming when presented in isolation, interference effects are found when the items are presented in a larger semantic contexts (e.g., in combination with other items related to a common theme; Abdel Rahman & Melinger, 2007; Rose & Abdel Rahman, 2016). Furthermore, context effects that are otherwise absent can be induced by meaningful cues highlighting semantic relations (Abdel Rahman & Melinger, 2010; Lin, Kuhlen, & Abdel Rahman, 2021). Based on these theoretical considerations and the available empirical evidence, we speculate that meaningful social contexts can bias planning processes towards more dominant facilitatory meaning processing.

In the current study we extended the classic PWI paradigm to a two-person setting embedded in a conversational turn sequence. One person produced the distractor word and the second person named the target picture as part of a simple card game in which two cards, distractor and target, had to be paired. Cards were either semantically related or unrelated. We manipulated the distance between the onset of the distractor word (as detected by a voice key) to the onset of the target (the appearance of the picture on the participant’s screen), the stimulus onset asynchrony (SOA), on two levels: In half of the trials the distractor preceded the target by 100ms (SOA-100), which typically produces strong semantic interference (e.g., Damian & Martin, 1999; Hantsch et al., 2009; Schriefers et al., 1990). For the other half of trials the distractor preceded the target by 650ms (SOA-650), which typically results in less interference, sometimes even facilitation, presumably because conceptual processing prevails (e.g., Alario et al., 2000; Bloem et al., 2004; Glaser & Düngelhoff, 1984). Experiment 2 implemented the identical experimental parameters in a classic single-subject PWI setting, using audio recordings of the distractors named spontaneously in Experiment 1. Experiment 3 tested a possible mechanism for reducing interference in communicative settings by changing parameters of the card game. In this variant of the card game participants were asked to assess the conceptual relationship between target and distractor, thus explicitly shifting their focus from lexical to conceptual representations.

We hypothesized that semantic interference typically observed in picture-word interference studies can in principle also be observed in communicative settings. Thus, utterances of one speaker may interfere with semantically related utterances of the next speaker. Furthermore we speculated that the degree of interference depends on the nature of the communicative situation. Specifically, the communicative game in Experiment 1 did not explicitly encourage participants to conceptually relate distractor and target to each other. Hence, participants may process the distractor word similarly as in non-communicative PWI implementations and experience semantic interference. On the other hand, the communicative setting itself may be sufficient to strengthen conceptual processing. If this is the case, semantic priming effects would be enhanced and diminish overall interference. The communicative game in Experiment 3 explicitly encouraged conceptual processing. Hence we expected a pronounced shift from semantic interference to facilitation. We tested these effects on two SOA levels.

Consistent with past studies, we expected semantic interference to be most pronounced at SOA-100ms. At SOA-650, semantic facilitation may be more pronounced. These hypotheses, the experimental procedures, and analyses were pre-registered (Kuhlen & Abdel Rahman, 2019). The raw data of the experiments reported here have been made available (Kuhlen & Abdel Rahman, 2020).

## Experiment 1

### Methods

#### Participants

Two native German speakers were recruited to each experimental session. Both served once in the role of the person naming the distractors and once as the person naming the target pictures. Participants were reimbursed, or received credit towards their curriculum requirements, and gave informed consent prior to participating in the study. Four participants had to be excluded and replaced because they had a laughing or coughing spell during the naming task. Furthermore, the audio recordings of twelve participants (naming the distractor words) turned out to be deficient and had to be replaced^1^ in order to have a complete set of recorded distractor stimuli for Experiment 2. The final analyses included 32 participants of a mean age of 26 (SD = 4.79), 23 women and 9 men.

Our sample size was determined prior to data collection through a power analysis that simulated the outcome of an LMM model testing for semantic interference at SOA-100ms (simr package, Green et al., 2016). The random structure of this model contained intercepts only and was specified on the basis of the first six participants (data of all further participants were simulated). We assumed an interference effect of 22.5ms, which corresponds in size to those found in previous PWI studies (e.g., Lorenz et al., 2018). With 32 participants we reached a power estimate of 85.8% chance (95% confidence interval: 83.48, 87.91) for detecting the hypothesized interference effect.

#### Materials

Our stimulus material consisted of 40 distractor words and 40 to be named color photographs depicting everyday objects, see Appendix A for a complete list of all object names. Pictures were scaled to 3.5cm x 3.5cm and presented on a homogenous grey background framed by an orange rectangle, resembling a playing card.

Pictures and words were combined to form semantically related and unrelated target-distractor pairings such that for each participant every picture and every word appeared in both conditions (semantically related and unrelated), and, in the same combination, in both SOAs. All target-distractor combinations were phonologically unrelated. Distractor objects were never target objects. Semantically related target-distractor pairs came from the same superordinate category (e.g., two types of birds, see supplementary materials). These semantically closely related objects have demonstrated strong interference effects in single-subject PWI experiments (Rose et al., 2019). Semantically unrelated target-distractor pairings were formed by re-combining the semantically related target-distractor pairs.

To reduce the predictability with which a particular target followed a given distractor, we created two distinct target-distractor sets within each semantic condition. A participant who was presented a certain set of distractor-target pairs when naming distractors would be presented with a different set when subsequently naming targets. The order in which these sets were presented was counter-balanced across teams. This procedure resulted in 80 related and 80 unrelated target-distractor pairs for each participant in the naming condition.

The presentation order of target-distractor pairs was pseudo-randomized using the software Shuffle_tk (Pallier, 2002) implementing the following constraints: (1) at least three trials separated the repetitions of a particular distractor word or target picture, (2) at least three trials separated the repetitions of a distractor word’s, or target picture’s, superordinate category. Under these constraints 64 stimulus lists were created, one for each participant and SOA condition.

Additionally, one semantically unrelated target-distractor pairing served as a first trial of each session and eight unrelated target-distractor pairings served as training trials. These did not enter data analyses.

#### Procedure

Two participants were invited per experimental session and consecutively served as Player D (naming the distractor) and Player T (naming the target), or vice versa. Upon arrival, participants were introduced to each other as team members. The card sorting task was explained to them using a set of laminated cards created from the training stimuli. The experimenter demonstrated the first round of the game, which begins with Player D asking the question “which card comes on”, then taking up a card from the stack and reading out the displayed word (e.g., “eagle”). Next, Player T draws a card and names the displayed picture (e.g., “vulture”). Then Player D places his card to the card corresponding to Player T’s picture. Participants played a few rounds among themselves, switching roles in between. To provide an incentive and to strengthen the team spirit the participants were told that the fastest and most accurate team would win a gift certificate of 25 EUR each. Participants were familiarized with the stimulus material, which was presented on paper as unsorted sets of pictures.

For the main experimental session, the two participants were seated in a sound- proof cabin sitting across from each other, each with a monitor and a microphone in front of them, see *Figure 1*. Trials began with a fixation cross. For Player D the fixation cross was replaced after 500ms with the written utterance “which card comes on…”, which was read aloud by Player D. After 1750ms the distractor word was presented, which was also read aloud by Player D. With the onset of the distractor naming, identified by a voice key, the target picture was presented with the given SOA (100 or 650ms later) on Player T’s monitor. Player T’s task was to name this picture as fast and accurately a possible. The onset of the target naming was recorded by a second voice key and resulted in our main dependent measure, picture naming latency. The onset of the target naming released the presentation of the decision screen on Player D’s monitor. The decision screen displayed two pictures side-by-side: the target picture that had been named by Player T and a filler picture that was not part of the stimulus set.

**Figure 1.**
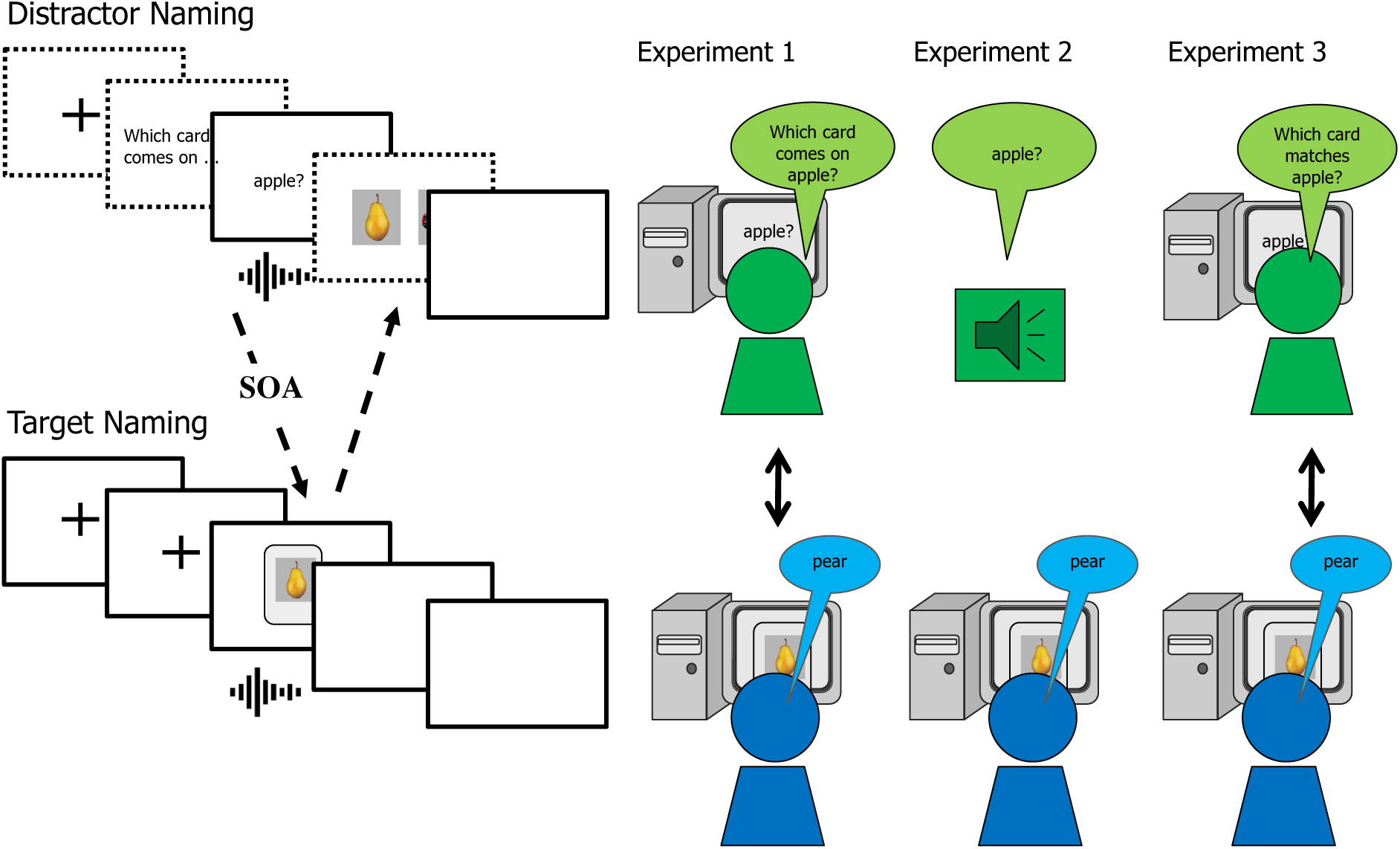
Illustration of trial structure and experimental setting of Experiments 1-3.

Player D indicated by button press where the distractor card should be placed. The target picture disappeared from Player T’s screen as soon as Player D gave a response. An Inter-Trial-Interval of 2000ms terminated each trial. Audio recordings were obtained from Player D’s reading of the distractor words (for details see Experiment 2).

SOA remained constant for one complete naming of all stimuli and then switched for the second round. The order of SOA condition was counter-balanced across participants. After playing out the entire set of stimuli in both SOA conditions, participants switched seats and roles. With swapped roles they played out all stimuli once more in both SOA conditions, but implementing different sets of target-distractor pairings. After completing both sessions the experimenter handed out a debriefing questionnaire.

#### Naming response

We analyzed the naming response of Player T, who named the target picture. Responses below 200ms were considered error trials and excluded from the analyses^2^ (4.67%). Furthermore excluded were trials during which no distractor-naming was registered or distractor naming was disfluent (1.97%) and trials in which target pictures were named wrongly or disfluently (4.14%). In total 91.21% of all trials were analyzed.

#### Data analyses

Linear mixed effects models (LMM; Baayen et al., 2008) as implemented in the *lmer* function of the *lme4* package (Bates, Maechler, Bolker, & Walker, 2015, Version 1.1-21) for R (R Development Core Team, 2014) were applied to log-transformed naming latencies. Naming latencies were modeled as a function of semantic relatedness (related vs. unrelated) in a nested structure, which tests for an effect of semantic relatedness at each SOA level (for model formula see Table 1). Both predictors were contrast coded, coding the first factor level as −0.5, and the second as +0.5.

**Table 1:**
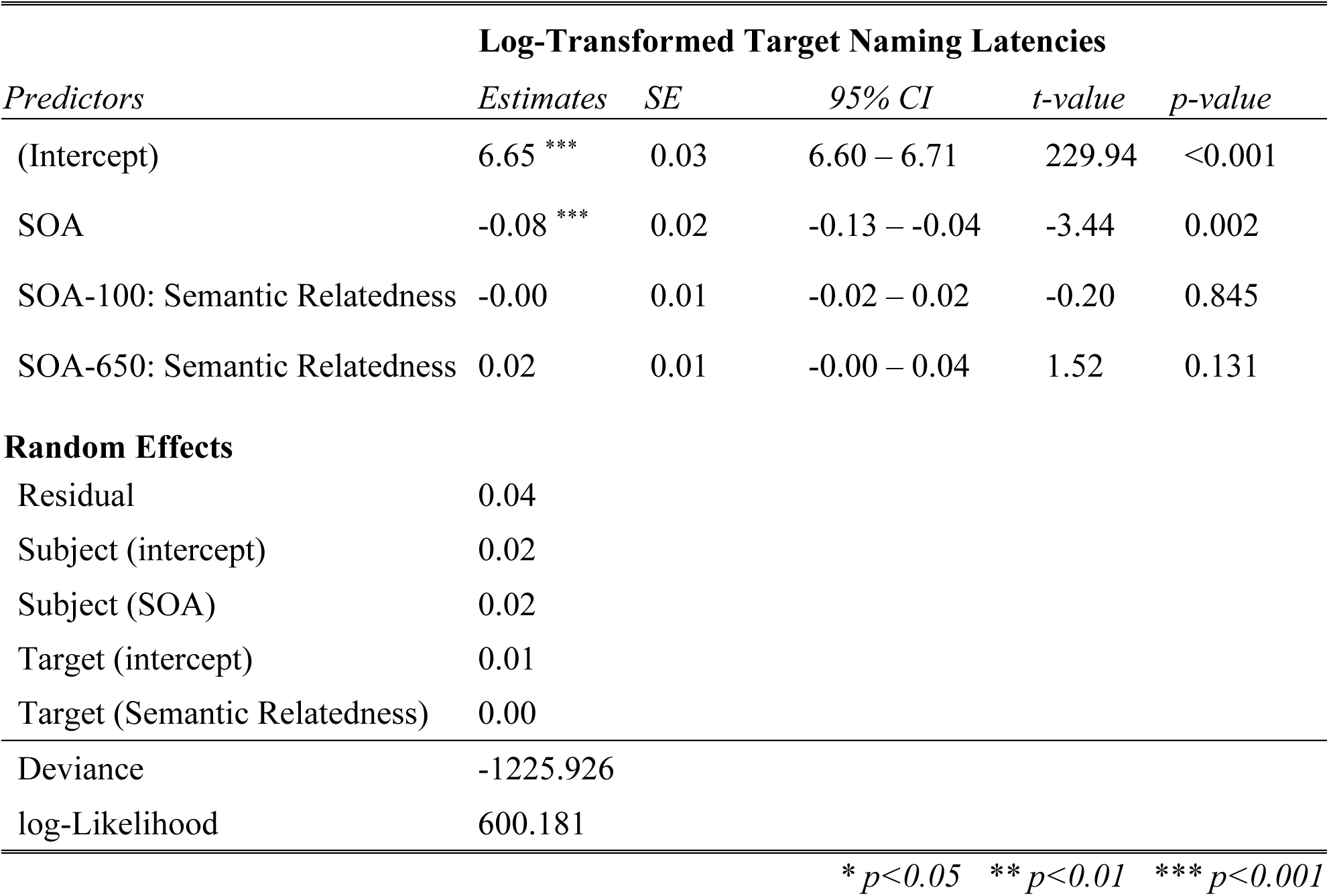
Model output for Experiment 1. Fixed-effect estimates, standard error, 95% confidence intervals, t-values, and p-values based on Satterthwaites approximations. Model equation: log(target naming latency) ∼ SOA/semantic relatedness + (SOA| subject) + (semantic relatedness| target).

Please note that we preregistered a model that tests an interaction between our predictors (semantic relatedness x SOA). However, as we realized during the analysis of Experiment 1, this model does not capture accurately our preregistered research question and hypotheses: primarily, we were interested in finding out whether semantic interference is experienced in a communicative setting. We expected semantic interference to be strongest at SOA-100. This effect is occluded when tested together with SOA-650. For SOA-650 predictions are less clearly derived from the literature and, if anything, are expected to go in the opposite direction. Thus, neither the predictor semantic relatedness alone nor an interaction between semantic relatedness and SOA capture well our research question. For our readers’ discretion we provide the models with the interaction term in Appendix C.

Models were initially run with a maximum random effects structure for participants and target pictures. Using singular value decomposition, the initial full random effect structure was simplified, if necessary, by successively removing those random effects for which estimated variance was indistinguishable from zero until the maximal informative model was identified (Bates et al., 2018).

For fixed effects, we report fixed effect estimates, standard errors, 95% confidence intervals, t-values, and p-values, as well as the square root (standard deviations) of the random effect structure, and goodness-of-fit statistics. P-values were computed via the Satterthwaite’s degrees of freedom method as implemented in the lmerTest package (Kuznetsova et al., 2017).

In preregistered control analyses we addressed the possible contribution of experimental design features: (1) the order in which participants played the role of the person naming the distractors (Player D) and the role of the person naming the targets (Player T), which captures the degree of familiarity participants have with the stimulus material at the time they name the targets, (2) the order in which participants experienced SOA-100 and SOA-650, which captures at each SOA level possible effects of naming the same target for the second time (practice effect), and (3) under which target-distractor pairing participants named targets. We tested the effect of these binary variables at each SOA level by entering them one at a time as additional predictors (contrast coded 0.5 vs. -0.5) to the identified final model, including an interaction with semantic relatedness.

To follow up on the pattern of our findings we conducted additional exploratory analyses including probing the stability of our models and exploring the distributional characteristics of semantic interference. These will be further motivated alongside the results.

### Results

#### Pre-registered hypotheses

Participants took on average 798.69ms (SD = 237.22) to name target pictures. Naming latencies were not affected by the semantic relationship between target and distractors: independent of whether their partner had just uttered a semantically related or unrelated word participants named target pictures at a comparable speed. This pattern could be observed for both short and long SOAs (mean difference related vs. unrelated condition at SOA-100 = 1.05, SE = 10.5; at SOA- 650 = -8.06, SE = 9.57). In both semantic conditions speakers named pictures faster after a long SOA of -650ms than after a short SOA of -100ms. The results are visualized in raincloud plots (modified from Allen et al., 2019) in *Figure 2*, the output of the LMM model can be found in *Table 1.* The maximal informative model included a random slope for SOA and a random slope for semantic relatedness.

**Figure 2.**
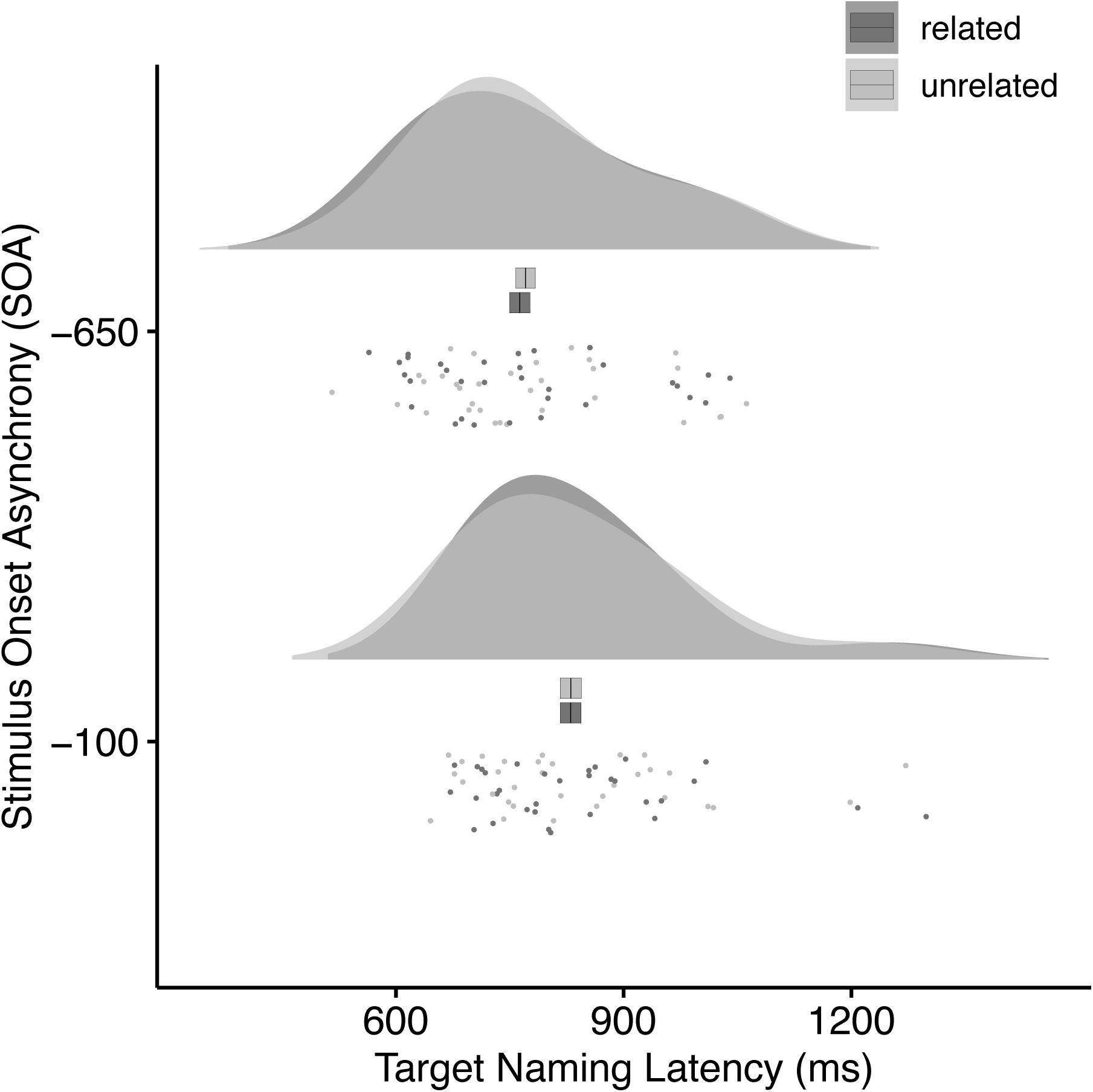
Data distribution of target naming latency (in milliseconds) as probability density function, including boxplots with mean values and 95% confidence intervals and individuals’ mean data points for each SOA and semantic relatedness condition separately of Experiment 1.

Our control analyses indicated that the design feature whether participants first named distractors and then targets, or vice-versa, influenced target naming latencies at both SOA levels: those participants who were already familiarized with the stimulus material through the prior iteration of the game named targets overall faster (SOA-100: β = -0.15, SE = 0.05, CI [-0.24 – 0.04], t = -2.98, p = 0.006; SOA-650: β = -0.14, SE = 0.05, CI [-0.25 – 0.04], t = -2.66, p = 0.013). On neither SOA level did this effect influence our main manipulation, semantic relatedness.

Relatedly, we observed that participants benefited from naming targets for the second time as the order under which participants experienced the short or long SOA affected target latencies: At SOA-100 participants named targets marginally faster when they had started with SOA-650 (β = -0.09, SE = 0.05, CI [-0.2 – 0.01], t = -1.72, p = 0.097) and at SOA-650 participants named targets significantly faster when they had started with SOA-100 (β = 0.15, SE = 0.05, CI [0.05 – 0.26], t = 2.8, p = 0.009). Thus, at both SOA levels naming was faster at the second round of naming. This practice effect did not influenced the effect of semantic relatedness.

Lastly, target naming latencies were not affected by the type of target-distractor pairing, indicating that in both sets of distractor-target pairings the effect of SOA and semantic relatedness played out similarly.

### Exploratory analyses

#### Analysis for overly influential cases

To address the possibility of individual participants or items overly influencing our model we re-estimated our model excluding individual participants and items using the ME.influence package (Nieuwenhuis, te Grotenhuis, & Pelzer, 2012). Figure D.1 in Appendix D visualizes the obtained dfbetas (Belsley, Kuh, & Welsch, 1980), a standardized measure of the absolute difference between the estimates specified by our original model as reported above and the estimates of a model excluding individual participants or items. As can be read off Figure D.1 six participants and three targets exceed the cut-off value recommended by Belsley and colleagues (1980) and were thus classified as having substantial influence on our estimates. Based on the recommendations of Nieuwenhuis and colleagues (2012) we subsequently re-ran our model excluding one influential case at a time. In eight of the nine cases this procedure did not change the outcome of our original model in a significant way. In one case, the exclusion of an individual participant resulted in a significant effect of semantic context at SOA-650 (Subject 14: β = 0.02, SE = 0.01, CI [0.00 – 0.04], t = 2.08, p = 0.041). This indicates that, without this participant, the remaining sample demonstrates a significant semantic facilitation effect at SOA-650. Removing the other influential participants or targets did not change the parameters of our original model in significant ways.

#### Exploration of distributional characteristics

Semantic interference in the picture-word interference task has been argued to be modulated by the degree of selective inhibitory control an individual engages in to suppress a semantically related distractor word (e.g., Shao et al., 2013, 2015). In the presence of another person participants may be particularly motivated to apply inhibitory control (see e.g., Sharma et al., 2010), thus attenuating semantic interference. To look into this possibility we visualized the observed naming latencies in vincentized cumulative distribution curves (e.g., Ratcliff, 1979; Roelofs, 2008). For these we rank ordered naming latencies for each subject and each condition separately and then divided these values into quintiles, see Figure E.1 of Appendix E. The first quintile in a given experimental condition reflects the mean value of the 20% of the trials with the fastest naming latency. Since selective inhibition builds up over time the effect of applying inhibition should be more pronounced on trials with slower naming latencies (e.g., Forstmann et al., 2008; Proctor, Miles, & Baroni, 2011; Ridderinkhof, 2002; Van den Wildenberg et al., 2011). Thus, semantic interference may have been present in trials with faster naming latencies, but this effect could have been masked by an increasing attenuation due to inhibitory control in trials with slower naming latencies. Yet if inhibitory control does not modulate our effect, interference would be more evenly distributed (Roelofs & Piai, 2017) or even be more pronounced for slower naming latencies, presumably because the distractor has more time to take effect (Ridderinkhof, 2002) or because slow naming latencies may indicate attentional glitches (Scaltritti et al., 2015). Since the effect should be strongest in short SOAs, so we extended our investigation to the SOA of -100ms only.

Based on the vincentized distribution plot, we find no indication that reduced semantic interference among the slower naming latencies may have been driving the attenuation of semantic interference overall. Instead, our data pattern shows an attenuated, but rather evenly distributed semantic interference effect across the middle and upper end of the naming latency distribution, a data pattern consistent with Roelofs and Piai (2017; see also Piai, Roelofs, & Schriefers, 2011). These observations are confirmed by an LMM analysis, in which we entered the obtained quintile bins as continuous predictor including an interaction with the predictor semantic relatedness. Our model indicates no interaction between the effect of semantic relatedness and quintiles (β = 0.00, SE = 0.01, CI [-0.01 – 0.01], t = 0.00, p = 0.997). We conclude that participants’ motivation to apply strong inhibitory control is unlikely to be driving the attenuation of semantic interference.

## Discussion

Based on the extensive literature on picture-word interference naming latencies should increase if the task partner produces a semantically related word in the previous turn. Especially at SOA-100 semantic interference is a well-documented and robust effect. Yet the results of Experiment 1 demonstrate that speakers do not experience semantic interference. We propose that the communicative game in which we embedded the picture-word interference paradigm prompted participants to relate the partner’s utterances and their own utterances conceptually to each other. This could lead to distractors priming targets and thus promote semantic facilitation and attenuate overall semantic interference (Abdel Rahman & Melinger, 2019, 2009). We report empirical support for this proposal in Experiment 3.

But could an alternative explanation be that the results of Experiment 1 were caused by unintended deviations from the basic PWI paradigm? Most notably, the SOA was implemented with a voice key and possible inaccuracies of the trigger (e.g., Kessler et al., 2002) could have resulted in irregular SOAs. Also, in the typical picture-word interference experiment auditory distractors are well-articulated and pre-recorded under optimal acoustic conditions. In our experiment, distractors were produced spontaneously and could have been less intelligible, and hence less effective. For these reasons we replicated the experiment with the audio recordings of the distractor naming obtained from Experiment 1 and implemented these with identical timing parameters in a single-subject setting devoid of a communicative context.

## Experiment 2

### Methods

#### Participants

For each experimental session, one native German speaker was invited to the lab at a time. Participants were reimbursed, or received credit towards their curriculum requirements, and gave informed consent prior to participating in the study. One participant had to be excluded due to non-compliance (repeated use of cell phone during the experiment). The final set of participants consisted of 32 people of a mean age of 27.84 years (SD = 5.26), 15 men and 17 women. None of the participants had participated in Experiment 1.

#### Materials

Stimulus materials were identical to those used in Experiment 1 with the exception that the distractor words consisted of all 32 audio recordings obtained from Experiment 1’s Player D. Audio recording started at the time point at which the written distractor had been presented to Player D and hence included a period of silence that varied depending on individual reading latency. The recordings ended with the completion of the word. Audio recordings were cleaned of static noise in Adobe Audition. The quality of all stimuli was checked manually. We marked as deficient recordings in which no distractor naming had occurred and recordings that contained speech disfluencies or recording artefacts that could not be corrected, in total 1.48% of all recordings. These deficient recordings were presented as stimuli, but excluded from the analyses. Recordings were presented via headphones to participants. The presentation order was identical to Experiment 1. This resulted in a session-by-session replication of the exact stimuli and their temporal characteristics as implemented in Experiment 1. What differed from Experiment 1 was the absence of a game partner and the lack of a communicative context.

#### Procedure

Participants were instructed to name the presented pictures as fast and accurately as possible. As is common in picture-word interference experiments, participants were told to ignore the distractor. To provide an incentive for fast and accurate responses comparable to Experiment 1 participants were told that the fastest and most accurate individual would win a gift certificate of 25 EUR. Prior to the main naming session, participants familiarized themselves with the stimulus material.

Each trial proceeded with the identical timing parameters as in Experiment 1: Trials began with a fixation cross presented for 500ms plus and additional 1750ms (the time period during which, in Experiment 1, Player D read their question). Then the audio recording began, implementing a voice onset of the distractor word identical to the one observed in Experiment 1. With the onset of the distractor word the target picture was presented on the participants’ monitor with the given SOA (100 or 650ms later). The onset of participants’ target naming was recorded by a voice key. The target picture disappeared from the participants’ monitor after a time period that corresponded to Experiment 1 (Player D’s decision where to place their card). An Inter-Trial-Interval of 2000ms terminated each trial. All other details of the design, presentation and data analysis were identical to Exp. 1

#### Naming response

Of all target naming responses 2.07% were excluded from the analyses because naming latency was below 200ms and 5.25% were excluded as error trials. Identical to Experiment 1, 1.97% of all trials were excluded due to missing or deficient distractor-naming. In total, 91.74% of all trials entered the analyses.

### Results

#### Pre-registered hypotheses

Participants named target pictures at a comparable speed as in Experiment 1, with an average naming latency of 793.22ms (SD = 241.74). At SOA-100 a pronounced interference effect was observed, which resembled in pattern and size those typically reported in PWI studies: participants took on average 22.21ms (SE = 10.98) longer to name the target picture when a semantically related versus unrelated distractor word had preceded it. Our LMM model confirmed this difference as statistically significant. The maximal informative model modeled variance across participants with a random slope for semantic relatedness and a random slope for SOA and variance across target pictures with a random slope for semantic relatedness.

At SOA-650 participants took on average 7.83ms (SE = 11.3) longer in the semantically related compared to the unrelated condition. This difference was not statistically significant. As in Experiment 1, participants were numerically faster to name target pictures after a long SOA (-650) than after a short SOA (-100), but this difference was not statistically significant. The results are visualized in *Figure 3*; the output of the LMM model is reported in *Table 2*.

**Figure 3.**
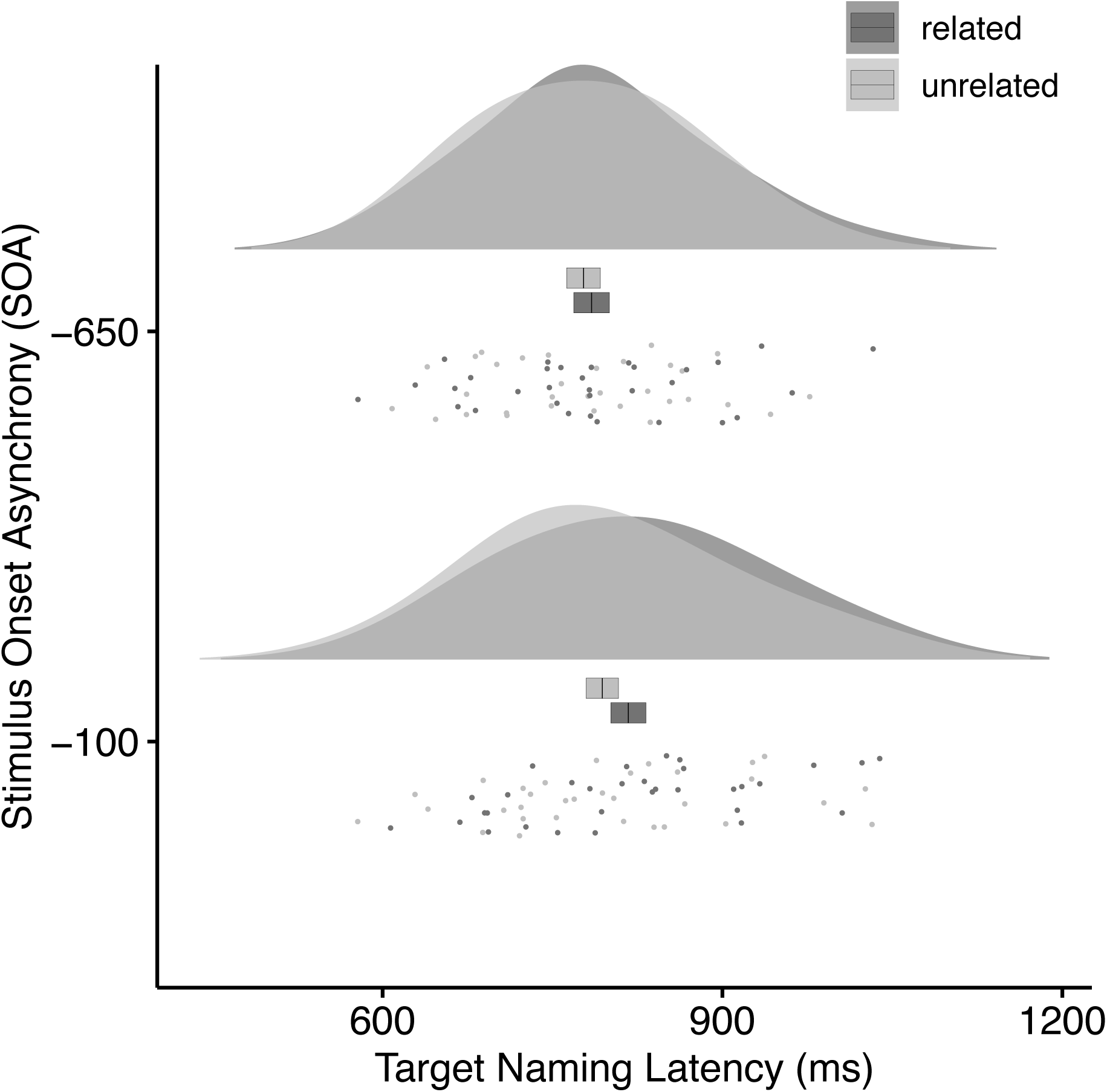
Data distribution of target naming latency (in milliseconds) as probability density function, including boxplots with mean values and 95% confidence intervals and individuals’ mean data points for each SOA and semantic relatedness condition separately of Experiment 2.

**Table 2:** Model output for Experiment 2. Fixed-effect estimates, standard error, 95% confidence intervals, t-values, and p-values based on Satterthwaites approximations. Model equation: log (target naming latency) ∼ SOA/semantic relatedness + (semantic relatedness+SOA| subject) + (semantic relatedness| target)

| **Log-Transformed Target Naming Latencies** |  |  |  |  |  |
| --- | --- | --- | --- | --- | --- |
| *Predictors* | *Estimates* | *SE* | *95% CI* | *t-value* | *p-value* |
| (Intercept) | 6.64 *** | 0.02 | 6.60 – 6.69 | 291.93 | <0.001 |
| SOA | -0.04 | 0.03 | -0.09 – 0.02 | -1.35 | 0.186 |
| SOA-100: Semantic Relatedness | -0.03 * | 0.01 | -0.05 – -0.00 | -2.24 | 0.029 |
| SOA-650: Semantic Relatedness | -0.01 | 0.01 | -0.03 – 0.02 | -0.43 | 0.667 |
| **Random Effects** |  |  |  |  |  |
| Residual | 0.05 |  |  |  |  |
| Subject (intercept) | 0.01 |  |  |  |  |
| Subject (Semantic Relatedness) | 0.00 |  |  |  |  |
| Subject (SOA) | 0.02 |  |  |  |  |
| Target (intercept) | 0.01 |  |  |  |  |
| Target (Semantic Relatedness) | 0.00 |  |  |  |  |
| Deviance | -226.582 |  |  |  |  |
| log-Likelihood | 100.631 |  |  |  |  |
| * $p < 0.05$ ** $p < 0.01$ *** $p < 0.001$ | | | | | |

Our control analyses indicated that, as in Experiment 1, participants’ target naming latencies decreased in their second naming of the target. This is indicated by an effect of SOA order: at SOA-100, naming latencies were faster when targets had already been named at SOA-650 (β = -0.16, SE = 0.04, CI [-0.24 – -0.09], t = -4.53, p < 0.001); and at SOA-650, naming latencies were faster when targets had already been named at SOA- 100 (β = 0.1, CI [0.02 – 0.17], SE = 0.04, t = 2.54, p = 0.017). This practice effect did not interact with the predictor semantic relatedness. The target-distractor pairing made no significant contribution to predicting naming latencies.

### Exploratory analyses

#### Analysis of overly influential cases

As in Experiment 1 we probed the stability of our findings by identifying possible participants or target items that may have overly influenced the estimates of our model (Nieuwenhuis et al., 2012). Two influential participants and five influential targets were identified, see Figure D.2 in Appendix D. Of these seven cases, the exclusion of two targets turned the semantic interference effect at SOA-100 to marginal significance (without Orang-Utan/orang utan: β = -0.02, SE = 0.01, CI [-0.05 – 0.00], t = -1.89, p = 0.063; without Pavian/baboon: β = -0.02, SE = 0.01, CI [-0.04 – 0.00], t = -1.91, p = 0.060). This suggests that these targets substantially strengthened the interference effect reported in this experiment. We conclude from these analyses that the classic semantic interference effect observed in Experiment 2 may be somewhat less precise, possibly due to the less controlled setting.

#### Effect of social conventions

One possible explanation for the strikingly different pattern we observe between Experiment 1 and Experiment 2 could lie in the social convention not to interrupt another speaker. Especially in a social setting, participants may have withheld their response thereby eliminating semantic interference. To look into this, we compared the data of Experiment 1 and 2 to investigate (a) whether a speaker’s naming latency depended on the duration of the distractor word (i.e., if the partner took longer completing the distractor word, did the speaker begin naming the target later?), and (b) whether the interturn interval between the end of the distractor word and the beginning of the target word was longer in Experiment 1 than in Experiment 2.

The word duration of each distractor was determined by calculating the root-mean-square of the audio signal’s amplitude and identifying the two most prominent changes in power, which typically represent the onset and offset of speech. Recordings for which only one change point could be found (0.8% of all files) and recordings for which the calculated word duration was below 250ms (4.08% of all files) were checked and manually corrected, if necessary. The average distractor word duration was 506.71ms (SD = 148.14). We set up an LMM which included as additional predictors experiment (contrast coded 0.5 vs. -0.5) and distractor duration (centered and entered as continuous variable). In both experiments distractor duration made a significant contribution to predicting naming latency. *Table F* in Appendix F reports the model’s output.

The average interturn interval was defined as the time period that elapsed between the offset of the distractor and the onset of the target naming. In Experiment 1 this interturn interval was on average 666.86ms (SD = 344.7), and in Experiment 2 659.97ms (SD = 364.36). This difference was not statistically significant, t (62) = 0.37, p = 0.71.

#### Exploration of distributional characteristics

In Experiment 1 we explored whether semantic interference may have been particularly attenuated at the tail end of the naming latency distribution, which could indicate inhibitory control mechanism as a possible driving factor. We did not find evidence for this. For comparison, we also investigated the distributional characteristics of the interference effect observed in Experiment 2. Visual inspection of the vincentized cumulative distribution curve (see *Figure E.2* in Appendix E) suggests that semantic interference is quite evenly distributed across the middle and (right) tail end of the distribution. Also our LMM analysis speaks for an even distribution of semantic interference as we found no significant interaction between semantic relatedness and quintile bins (β = -0.01, SE = 0.01, CI [-0.02 – 0.01], t = -0.90, p = 0.373). This broad distribution of semantic interference is consistent with the pattern reported by Roelofs and Piai (2017).

### Discussion

A session-by-session replication of Experiment 1 that did not elicit semantic interference in a social setting did elicit semantic interference when embedded in a single-subject setting. The pattern of results at SOA-100 resembles the typical finding in PWI studies. We do not find any influence of semantic relatedness at SOA-650, at which semantic effects are considerably less stable. Taken together, we conclude that our specific implementation of the PWI paradigm is not responsible for diminishing semantic interference in Experiment 1.

In dialogue it is typically considered inappropriate to begin speaking before the previous speaker has completed their turn (e.g., Sacks et al., 1974). This convention may have occluded semantic effects in Experiment 1. Yet our exploratory analyses suggests that similarly in a single-subject setting participants may be withholding their response while the distractor is being articulated. This makes it unlikely that semantic interference was reduced because participants delayed their response in a social setting.

Instead we suggest that the game in which the PWI paradigm was embedded in Experiment 1 (but not in Experiment 2) encouraged participants to process the partner’s utterances more strongly on a conceptual level. When conceptual processing is emphasized, semantic relatedness has been suggested to alleviate interference, and perhaps even turn to facilitation (e.g., Abdel Rahman & Melinger, 2019). To test this idea, we implemented in Experiment 3 a modified communicative game designed to increase participants’ focus on the conceptual commonalities between distractor and target. Resulting facilitatory effects are most likely to be found at SOA-650.

## Experiment 3

### Methods

#### Participants

As in Experiment 1 we recruited two native German speakers per session. All participants gave informed consent prior to participating in the study and were reimbursed or received credit towards their curriculum requirements for their participation. Two participants had to be excluded due to laughing spells. The final set of 32 participants had a mean age of 22.91 years (SD = 4.59), 4 men and 28 women.

#### Materials

The stimulus materials and lists were identical to those employed in the first two experiments.

#### Procedure

The experiment proceeded as Experiment 1 with the following changes to the communicative game: The game began with Player D asking the question “What matches…” (instead of the semantically less relevant “Which card comes on…”) followed by the naming of the distractor word. With a given SOA added to the onset of the distractor word, the target picture appeared on Player T’s monitor, who then named the displayed object as fast and accurately as possible. At the end of each trial Player D indicated whether the picture named by Player T matched the distractor word. If participants were uncertain what counted as a matching card the experimenter gave an example (“Piano and violin match, they are both instruments.”). As in the previous experiments, the fastest and most accurate team won a prize. Naming responses and SOAs were determined as in the previous experiments. All other details of the procedure and data analysis were identical.

#### Naming response

As in Experiment 1-2 we analyzed the naming response of the participants naming the target picture. 89.9% of all trials entered the analyses. Excluded were 1.23% due to missing distractor naming, 6.41% due to wrong or disfluent target naming, and 3.71% due to naming latencies below 200ms.

### Results

#### Pre-registered analyses

Participants named target pictures with an average naming latency of 845.66ms (SD= 227.14). The data pattern replicated the pattern observed in Experiment 1 for SOA-100: the typically observed semantic interference effect was greatly diminished and did not reach statistical significance (mean difference related vs. unrelated = 10.26ms, SE = 10.3). Instead, at SOA-650ms, we observed a statistically significant facilitatory effect of semantic relatedness: after their partner uttered a semantically related distractor word speakers were on average 30.15ms (SE = 14.03) faster to name the target picture. The results are visualized in *Figure 4*; the output of the model is reported in *Table 3*. Our data supported a model accounting for variance across participants with a random slope for semantic relatedness and SOA and across target pictures with a random slope for semantic relatedness and SOA.

**Figure 4.**
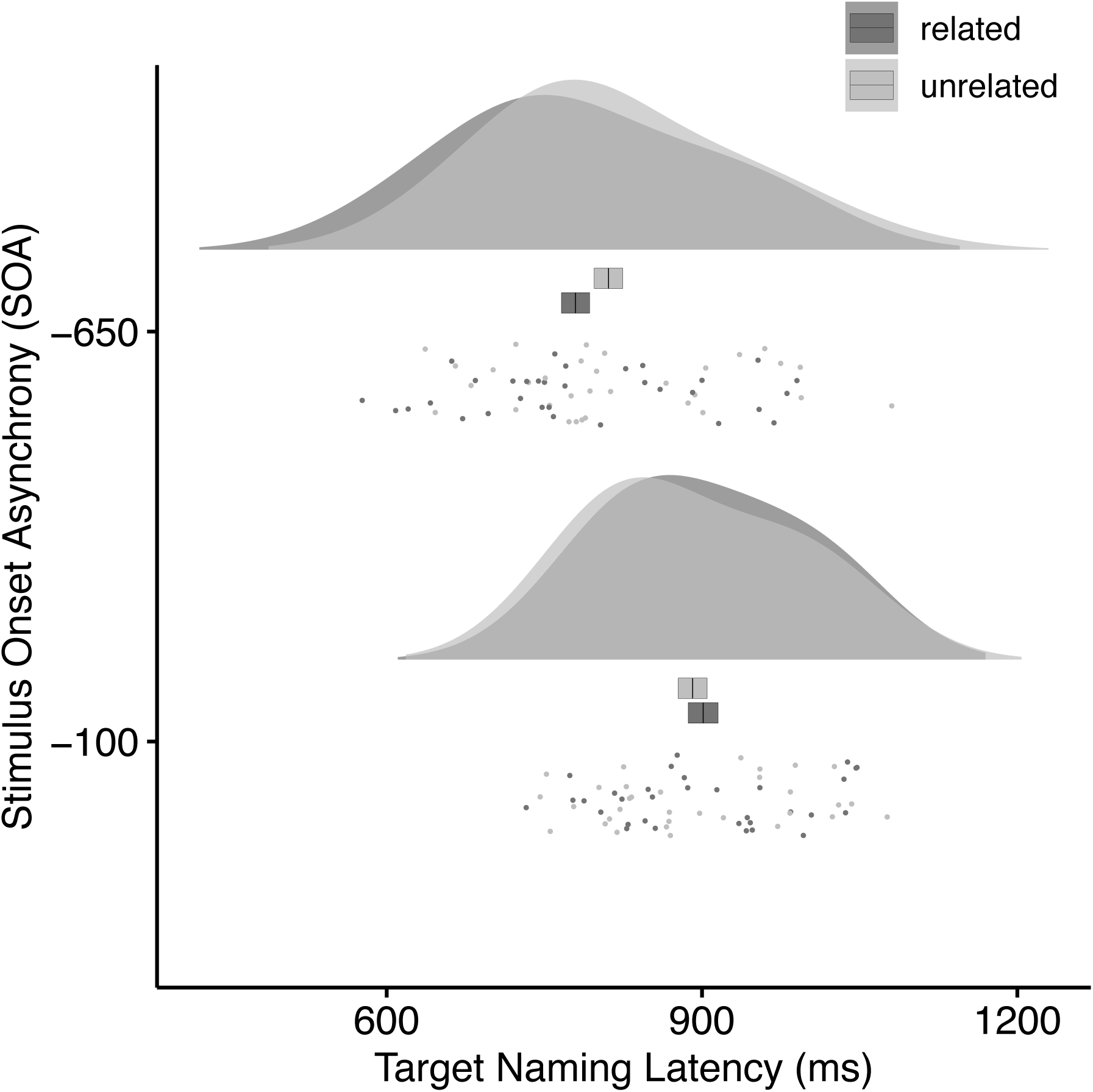
Data distribution of target naming latency (in milliseconds) as probability density function, including boxplots with mean values and 95% confidence intervals and individuals’ mean data points for each SOA and semantic relatedness condition separately of Experiment 3.

**Table 3:**
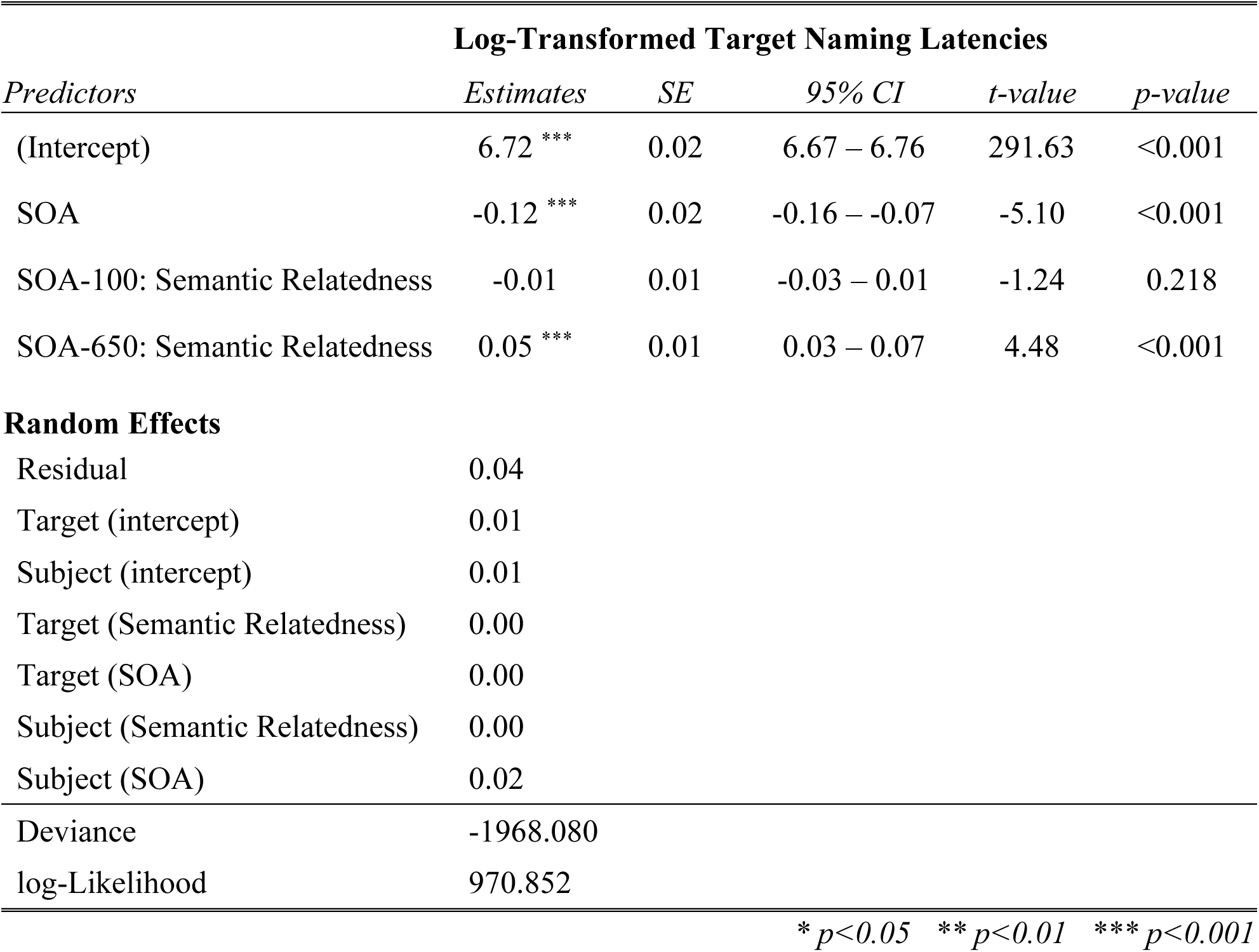
Model output for Experiment 3. Fixed-effect estimates, standard error, 95% confidence intervals, t-values, and p-values based on Satterthwaites approximations. Model equation: log(target naming latency) ∼ SOA/semantic relatedness + (semantic relatedness+SOA| subject) + (semantic relatedness+SOA| target)

Our control analyses indicated that, as in Experiment 1, naming latencies decreased as participants became familiar with the stimulus material: Participants named targets faster when their partner had named the targets in the prior iteration of the game (vs. when participants were first to name targets). This yielded a statistically significant effect at SOA-100 (β = -0.1, SE = 0.03, CI [-0.16 – -0.04], t = -3.17, p = 0.004), and was numerically, yet not statistically, confirmed at SOA-650 (β = -0.07, SE = 0.05, CI [-0.16 – 0.02], t = -1.49, p = 0.148). This familiarity effect did not influence the effect of our main manipulation, semantic relatedness. Likewise, as participants named targets for the second time their naming latencies decreased, as indicated by an effect of SOA order, numerically at SOA-100 (β = -0.04, SE = 0.04, CI [-0.11 – -0.03], t = -1.08, p = .288) and statistically significant at SOA-650 (β = 0.19, SE = 0.03, CI [0.12 – 0.25], t = 5.52, p < 0.001). At both SOA levels, this practice effect seemed to add on top of the effect of semantic relatedness: At SOA-100 semantic interference was marginally smaller when targets were named for the second time (β = 0.03, SE = 0.02, CI [-0.00 – 0.06], t = 1.74, p = 0.085); at SOA-650 semantic facilitation was significantly larger when targets were named for the second time (β = -0.05, SE = 0.02, CI [-0.08 – -0.01], t = -2.77, p = 0.007). At both SOA levels this did not change the pattern of our main findings: regardless of whether participants named targets for the first or for the second time, overall no semantic context effect emerged at SOA-100 (β = -0.01, SE = 0.01, CI [-0.03 – -0.01], t = - 1.25, p = 0.216), and a pronounced semantic facilitation effect emerged at SOA-650 (β = 0.04, SE = 0.01, CI [0.02 – 0.06], t = -4.35, p < 0.001). As in the previous experiments, the specific target-distractor pairing experienced by a given participants did not contribute to predicting target naming latencies.

### Exploratory analyses

#### Analysis of overly influential cases

We identified three participants and seven influential targets that had a significant influence on our model. For neither of these cases did an exclusion change the pattern of our findings in a significant way. We conclude that these cases substantially contribute to our findings but do not alter the conclusion we reached from our original model.

#### Exploration of distributional characteristics

To look into distributional characteristics of our effects, we computed vincentized cumulative distribution curves, see *Figure E.3* in Appendix E. These indicate that semantic interference may be slightly more prominent among slower naming latencies, a pattern that is inconsistent with inhibitory control attenuating semantic interference. An interaction of semantic relatedness and quintile bins was not statistically confirmed in our LMM model (β = 0.0, SE = 0.01, CI [-0.01 – 0.01], t = 0.18, p = 0.857). We conclude that, similar as in Experiment 1, pronounced inhibitory control in social settings is unlikely to be driving the observed attenuation of semantic interference.

#### Combined analyses Experiments 1-3

We combined the data from all three experiments and set up an LMM analysis contrasting each social setting with the single subject setting (Experiment 1 vs. Experiment 2; Experiment 3 vs. Experiment 2). We were most interested in probing our two main findings, the diminished semantic interference in a social setting at SOA-100 and the presence of semantic facilitation at SOA-650 in Experiment 3. Indeed, the effect of semantic relatedness differed between Experiment 1 and 2 at SOA-100, confirming our observation that semantic interference is significantly reduced in a social setting. However, this pattern did not reach statistical significance when comparing Experiment 3 with Experiment 2. At SOA-650, semantic relatedness had a distinct effect at SOA-650 in Experiment 3 compared to Experiment 2. This confirms the strong semantic facilitation we observe at this SOA in a social setting. For the full model output please refer to *Table 4*.

**Table 4:** Model output for global LMM pooling data from Experiment 1, 2 & 3. Fixed-effect estimates, standard error, 95% confidence intervals, t-values, and p-values based on Satterthwaites approximations. Model equation: log(target naming latency) ∼ Experiment * (SOA/semantic relatedness) + (SOA+semantic relatedness| subject) + (SOA+semantic relatedness| target).

| *Predictors* | **Log-Transformed Target Naming Latencies** |  |  |  |  |
| --- | --- | --- | --- | --- | --- |
|  | *Estimates* | *SE* | *CI* | *t-value* | *p-value* |
| (Intercept) | 6.67 *** | 0.02 | 6.63 – 6.71 | 363.00 | <0.001 |
| Exp2-1 | -0.01 | 0.03 | -0.07 – 0.05 | -0.33 | 0.744 |
| Exp3-2 | 0.07 * | 0.03 | 0.02 – 0.13 | 2.48 | 0.015 |
| SOA | -0.08 *** | 0.01 | -0.11 – -0.05 | -5.48 | <0.001 |
| SOA-100:Semantic Relatedness | -0.01 | 0.01 | -0.03 – 0.00 | -1.65 | 0.103 |
| SOA-650:Semantic Relatedness | 0.02 * | 0.01 | 0.00 – 0.03 | 2.27 | 0.027 |
| Exp2-1:SOA | 0.05 | 0.04 | -0.02 – 0.12 | 1.39 | 0.167 |
| Exp3-2:SOA | -0.08 * | 0.04 | -0.15 – -0.01 | -2.36 | 0.020 |
| Exp2-1:SOA-100:Semantic Relatedness | -0.03 * | 0.01 | -0.05 – -0.00 | -2.01 | 0.045 |
| Exp3-2:SOA-100:Semantic Relatedness | 0.01 | 0.01 | -0.01 – 0.04 | 1.08 | 0.282 |
| Exp2-1:SOA-650:Semantic Relatedness | -0.02 | 0.01 | -0.05 – 0.00 | -1.62 | 0.106 |
| Exp3-2:SOA-650:Semantic Relatedness | 0.05 *** | 0.01 | 0.02 – 0.07 | 3.94 | <0.001 |
| **Random Effects** |  |  |  |  |  |
| Residual | 0.04 |  |  |  |  |
| Subject (intercept) | 0.01 |  |  |  |  |
| Subject (SOA) | 0.02 |  |  |  |  |
| Subject (Semantic Relatedness) | 0.00 |  |  |  |  |
| Target (intercept) | 0.01 |  |  |  |  |
| Target (SOA) | 0.00 |  |  |  |  |
| Target (Semantic Relatedness) | 0.00 |  |  |  |  |
| Deviance | -3377.608 |  |  |  |  |
| log-Likelihood | 1649.808 |  |  |  |  |

\* $p < 0.05$ \*\* $p < 0.01$ \*\*\* $p < 0.001$

### Discussion

The communicative game of Experiment 3 was designed to prompt participants to evaluate the conceptual relationship between the partner’s and their own utterance. And, indeed, in this setting we observed semantic facilitation at SOA-650, a time window during which conceptual processing, and hence semantic priming, typically dominates. As we did not find facilitatory effects at this SOA in our previous experiments this finding supports our proposal that conceptual processing was promoted by the nature of the communicative game.

Moreover, Experiment 3 replicated our initial finding of greatly reduced semantic interference in communicative context at SOA-100. While in Experiment 1 we could not detect semantic interference even numerically, in Experiment 3 we observed a weak numeric semantic interference effect. Hence, when directly compared to our single-subject version of the PWI paradigm, the reduction of the classic interference effect is less pronounced as in Experiment 1. This may be connected to the overall longer naming latencies observed in Experiment 3. Longer naming times may strengthen interference effects (e.g., Bürki et al., 2020; Scaltritti et al., 2015). However note that our distributional analyses suggest that interference effects are not exclusively found in longer naming latency. In sum, both experiments show converging data patterns: a pronounced attenuation of semantic interference at a time window during which semantic interference should dominate according to traditional picture-word interference experiments. We will discuss the implications of these findings below.

## General Discussion

Traditional accounts of speech production are largely based on tightly controlled experimental settings in which a single subject is instructed to name pictures devoid of a communicative purpose. In these settings it has been demonstrated that speaking is delayed when a semantically related word is processed in close temporal proximity to the target picture (picture-word interference). Yet in dialogue interlocutors often refer to semantically related concepts across speaking turns. We sought out to investigate whether semantic interference can be observed when a word uttered in one task partner’s turn semantically relates to a word uttered in the other task partner’s turn. For this we implemented a classic picture-word interference paradigm in a communicative setting. In two experiments we did not observe semantic interference. Yet the same experimental design and stimuli, implemented in a single-subject setting, did yield the typical semantic interference effect. We conclude that in dialogue-like settings semantic interference elicited by processing and producing semantically related words in close temporal proximity is greatly diminished.

Our findings are in line with a recent publication that embedded the PWI paradigm in a discourse context (Shao & Rommers, 2019, Experiment 1). In this experiment participants heard a question, pre-recorded on audio, which was either tightly or loosely linked to the target picture. Immediately after the question, the target picture was presented alongside a semantically related or unrelated written distractor. When the question constrained as an answer the target picture, interference by a semantically related distractor was reduced (see e.g., Kleinman et al., 2015 for comparable findings). The authors conclude that discourse context can facilitate word selection by constraining semantic processing. Our study differs from this study in decisive ways: In Shao and Rommers’ study the distractor word intruded the communicative context by giving additional, misleading context. In our study, the distractor was part of the communicative context established by the task partner’s turn. And even though the partner’s utterance did not constrain the speaker’s naming response (current Experiment 1) we found no semantic interference. Thus, the driving forces behind the attenuation of semantic interference reported in Shao and Rommers’ study and in our present study may differ.

Our novel findings can be explained by models assuming that semantic context can have both facilitatory and interfering effects on speech production (e.g., Roelofs, 2018). Such models allow for opposing effects of identical semantic relations depending on whether the context or task at hand promotes conceptual processing (eliciting facilitatory priming) or lexical processing (eliciting competition) (Abdel Rahman & Melinger, 2009; 2019). Indeed, our results suggest that the communicative context we established in our modified PWI paradigm promotes processing the conceptual relationship between the partner’s and one’s own utterances. This results in the distractor, named by the partner, priming the target concept and its affiliated lexical representation and thus facilitating target selection. Hence, facilitatory priming on a conceptual level and interference on a lexical level canceled each other out (Experiment 1), or even turned towards facilitation when the communicative context explicitly emphasized the semantic relation between target and distractor (Experiment 3). A conceptual locus is also in line with the finding that facilitation was found when the word was presented 650ms before the picture.

Alternative accounts of word production that do not assume a trade-off between lexical competition and conceptual priming have been proposed to explain semantic interference effects in picture-word interference paradigms (e.g., Finkbeiner & Caramazza, 2006; Mahon et al., 2007). According to these accounts interference between target and competitor arises after lexical selection, at the level of articulation. Since only one lexical entry can pass through the so-called articulatory output buffer at a time, distractor and target impede each other during articulation. This clog resolves more quickly if the competing item is not response relevant. According to these original proposals, response relevance is defined by the degree of semantic relatedness between a distractor and a target word. If the definition of response relevance were to be extended to include a higher-level assessment of the contextual or pragmatic relevance at large, one could argue that a distractor word uttered as part of a conversational partner’s turn may not be considered response relevant and hence may elicit less interference. However, recent studies speak against a constraining influence of pragmatic context on word selection (Jescheniak et al., 2017; Mädebach et al., 2020).

A second alternative explanation for our findings comes from studies that have investigate interference effects in joint actions (e.g., Sharma et al., 2010; Sellaro et al., 2018). Most notably, a recent conference contribution (Tufft & Richardson, 2020) reports a picture-word interference experiment in which, comparable to our study, the naming of the written distractor word and the naming of the target picture is distributed between two task partners. In this study, the naming of the distractor and target is devoid of a communicative purpose. As in our study, semantic interference disappears. The authors suggest that the shared task setting prompts participants to off-load the processing of the distractor to the partner (interestingly, even if the partner’s task does not involve naming the distractor, but instead naming the color of the frame on which the picture-word pair is presented). In our study, as well, the shared task context may have alleviated participants from processing the distractor word deeply. Yet, social off- loading cannot explain why changes to the communicative game, most notably an increased focus on the conceptual relationship between distractor and target, would facilitate speech production.

Interestingly, in Tufft and Richardson’s study the interference effects becomes increasingly smaller with longer reaction times—a pattern of result that is consistent with the exertion of late inhibitory control. In our data we do not find an indication that participants were exerting inhibitory control to suppresses the processing of the distractor. This is a further indicator that there may be different mechanisms at play that can reduce interference in social settings. This will be a fruitful field for future research.

A related question concerns the level of detail at which a task partner’s utterance is represented. This is a controversial topic among recent studies that have implemented different joint picture naming paradigms to investigate speech production in shared task setting. Some of these studies demonstrate that pictures that are (presumably or actually) named by a partner contribute to the degree of interference a partner experiences, suggesting that speakers engage in lexical access on behalf of their partner (Baus et al., 2014; Hoedemaker et al., 2017; Kuhlen & Abdel Rahman, 2017). Yet, other studies have reported that speakers may not represent their partner’s naming response, or at least not at the lexical level (Brehm et al., 2019; Gambi et al., 2015; Hoedemaker & Meyer, 2019; Tufft & Richardson, 2020). Based on the greatly diminished semantic interference our current findings could be taken as indication that the partner’s utterances are not, or at least not predominantly, represented on the lexical level. Given the facilitatory effects observed in Experiment 3, our results however do not support the conclusion that the partner’s utterances were not represented at all. While previous studies investigated speech production in settings involving a second speaker, these settings were not, as in the current study was, embedded in a communicative exchange. The divergent findings across different experimental paradigms and social settings highlight the need for more systematic investigations of the processes underlying speech production in social interaction.

In dialogue, interlocutors typically speak about a particular topic. In doing so they are likely to refer to semantically related concepts across speaking turns. This could involve speaking about different members of a given semantic category (e.g., when discussing the menu options at a restaurant) or it could also involve referring to concepts that share a thematic relationships (e.g., are tied to a restaurant visit). Both types of semantic relationships have been shown to elicit semantic interference during language production (see e.g., Abdel Rahman & Melinger, 2007, 2010; Rose & Abdel Rahman, 2016). While communicative situations are typical instances for settings in which meaning-based processing may dominate, and hence reduce semantic interference, we do not assume that communicative situations are unique. Instead, our findings point towards a highly flexible semantic system that adapts to a broad range of semantic contexts and goals.

In sum, our results demonstrate that semantic context is processed differently when it is embedded in a communicative exchange compared to when it is embedded in a setting devoid of a social context (i.e., the typical PWI paradigm). Our findings suggest that a communicative setting promotes processing the conceptual relationship between distractor and target and hence enhances semantic priming. Lastly, our findings highlight the relevance of investigating language production in settings in which it typically occurs, namely in social interaction.

## Authors’ Contribution

Both authors contributed to the study design. Testing and data collection were performed by student assistants. A. K. Kuhlen performed the data analysis and interpretation with significant contributions by R. Abdel Rahman. A. K. Kuhlen drafted the manuscript, and A. Abdel Rahman provided critical revisions. Both authors approved the final version of the manuscript for submission.

## Acknowledgements

This research was supported by grants AB277/11-1 and AK3236/3-1 from the German Research Council to Rasha Abdel Rahman and Anna K. Kuhlen. We thank Jolanda Pogade, Nora Holtz, and Annika Just for assistance in data collection and analyses, and Guido Kiecker for technical support.

## Appendix A

**Table A:**
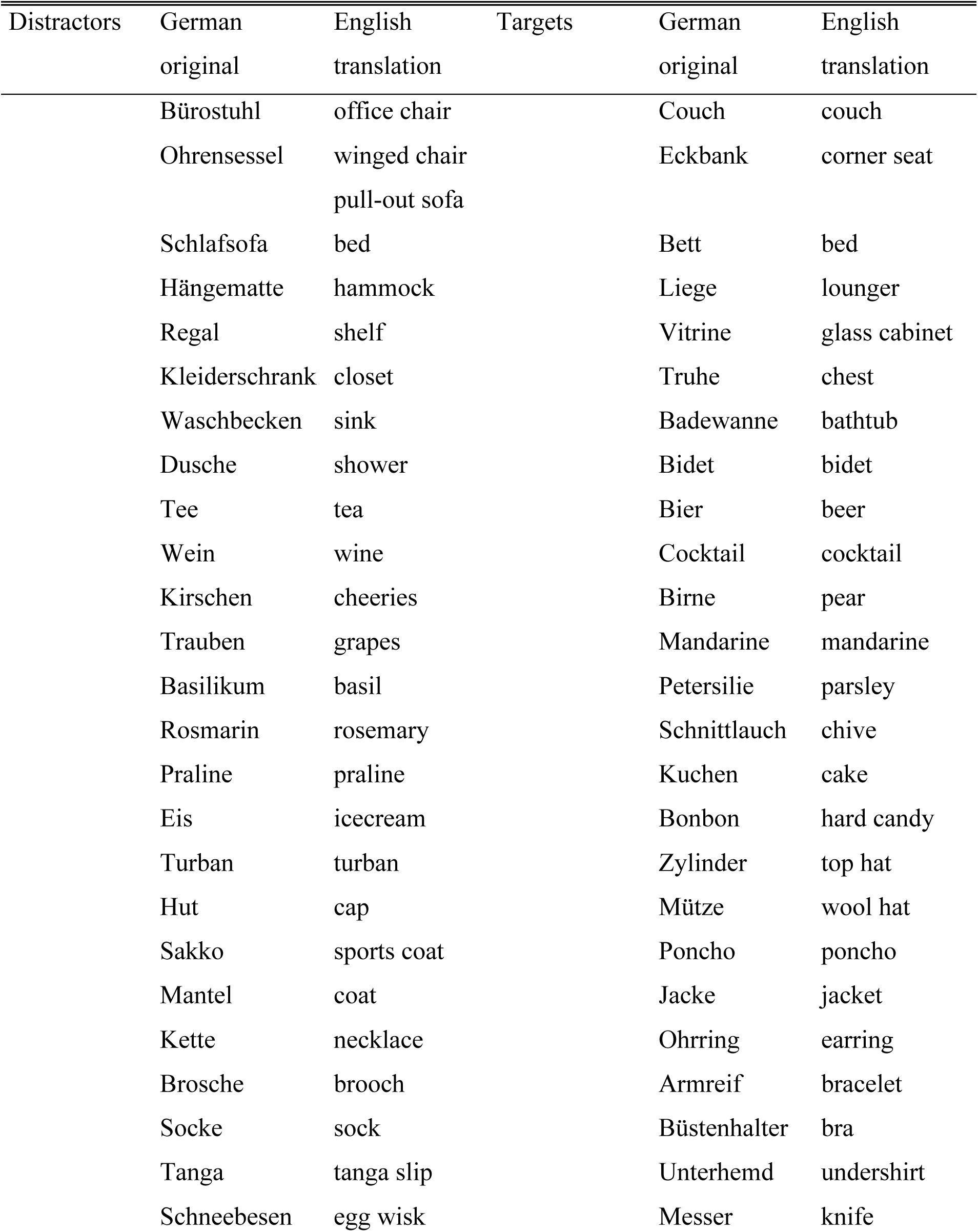
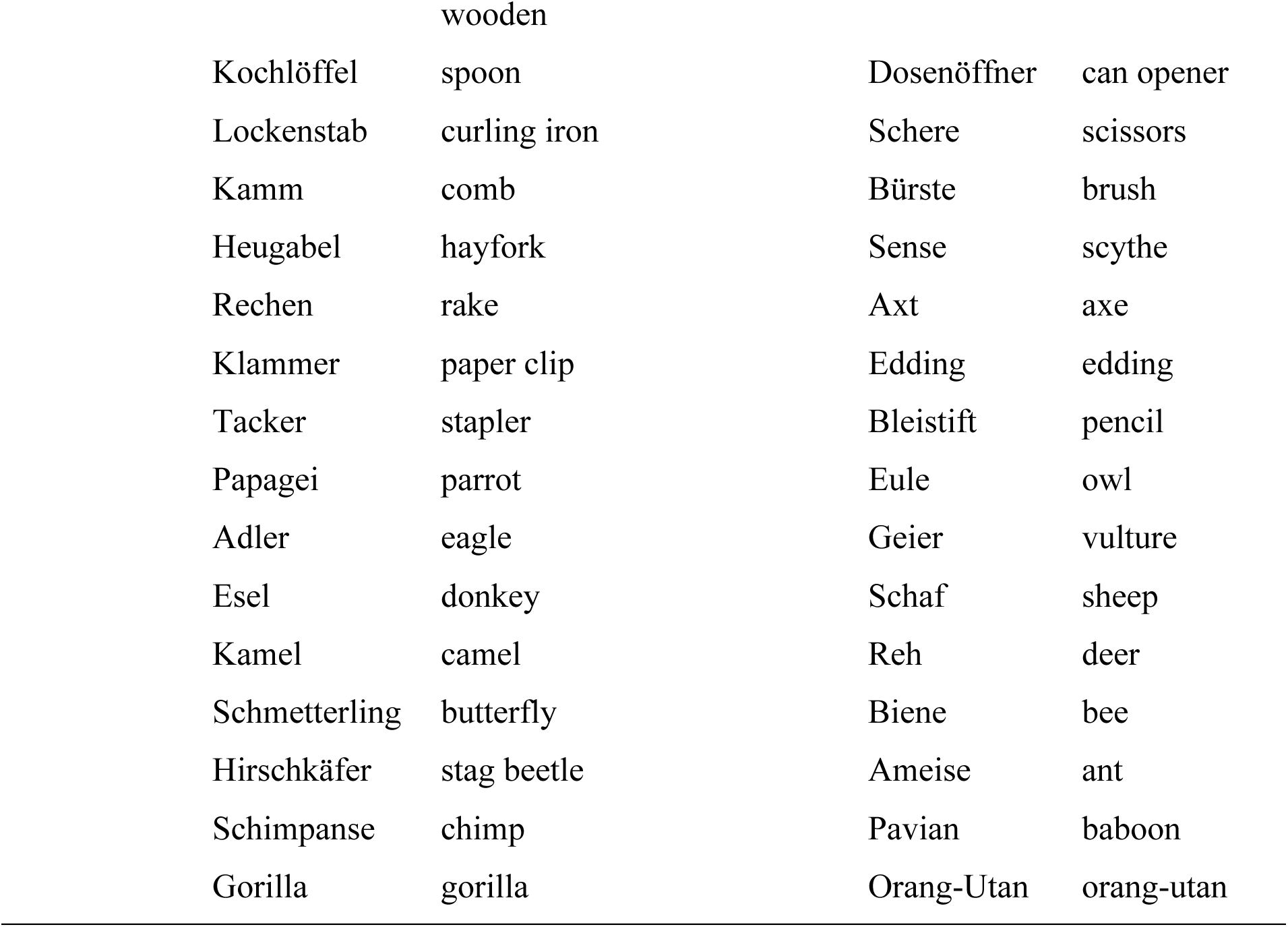
Complete list of object names used as distractors and targets respectively. Each line represents a semantically related target and distractor pairing. A second semantically related pairing was created by recombining target and distractors of the line below. Unrelated pairings were achieved by recombining targets and distractors of different semantic categories. Distractors were presented as written words to be read by the first speaker, targets were presented as pictures to be named by the second speaker.

## Appendix B

**Figure B.1.**
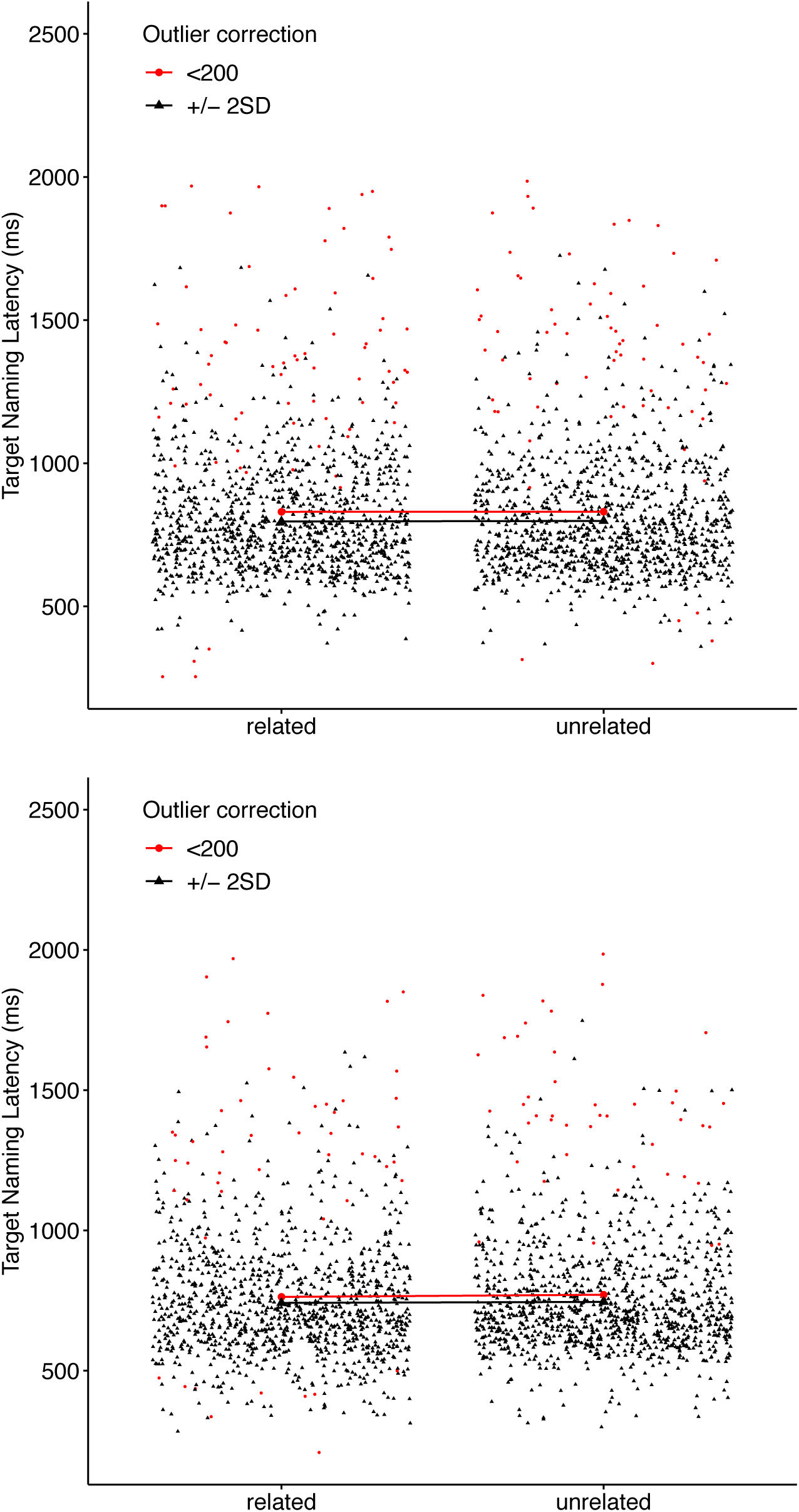
Experiment 1’s raw target naming latencies for each subject and item in the semantically related and unrelated condition for SOA-100 (top panel) and SOA-650 (bottom panel). Items marked as red circles would be excluded based on the pre- registered outlier correction (excluding trials +/-2SD of an individual’s mean). The implemented outlier correction (excluding trials <200ms) is less invasive.

**Figure B.2.**
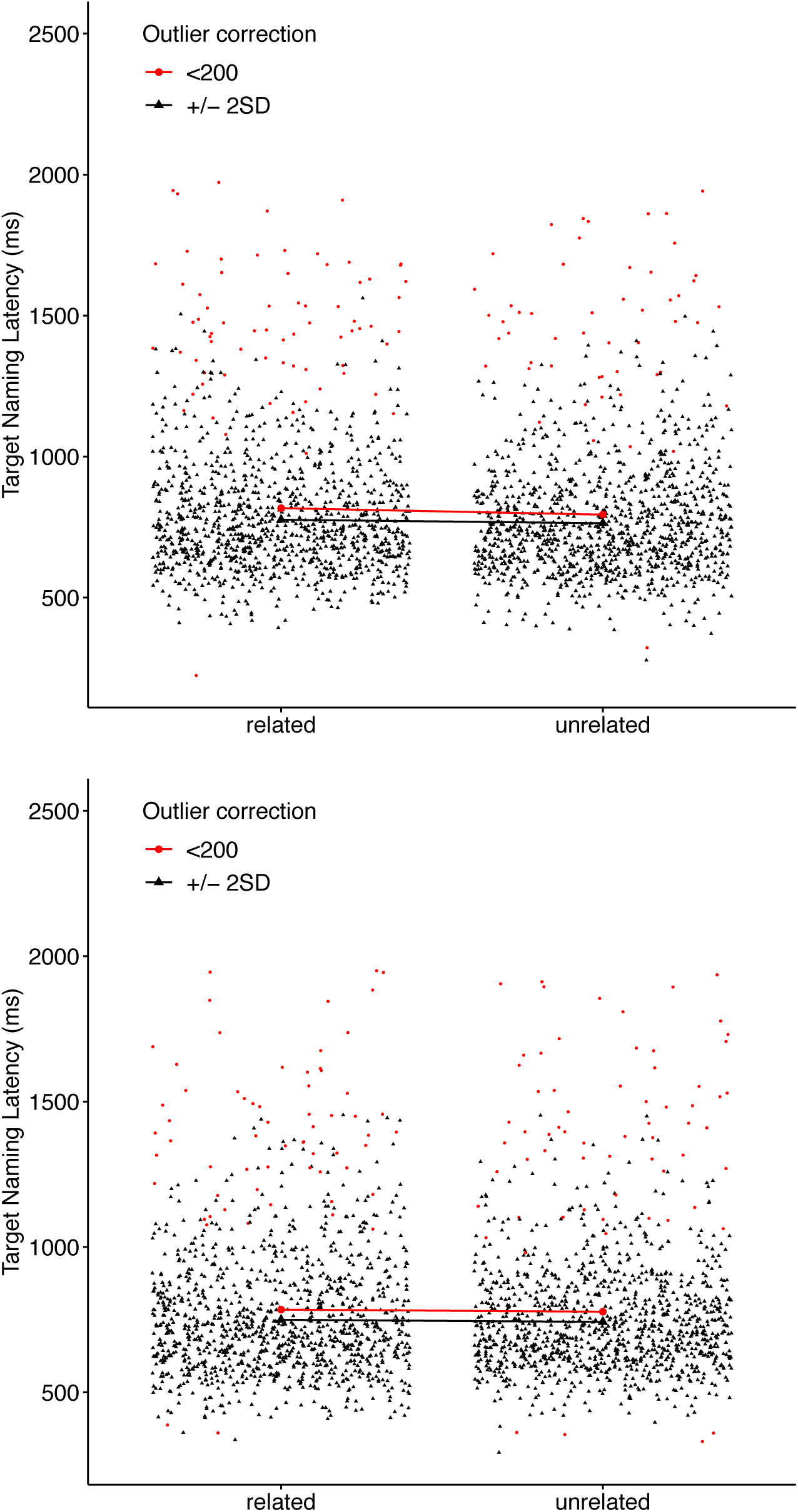
Experiment 2’s raw target naming latencies for each subject and item in the semantically related and unrelated condition for SOA-100 (top panel) and SOA-650 (bottom panel). Items marked as red circles would be excluded based on the pre- registered outlier correction (excluding trials +/-2SD of an individual’s mean). The implemented outlier correction (excluding trials <200ms) is less invasive.

**Figure B.3.**
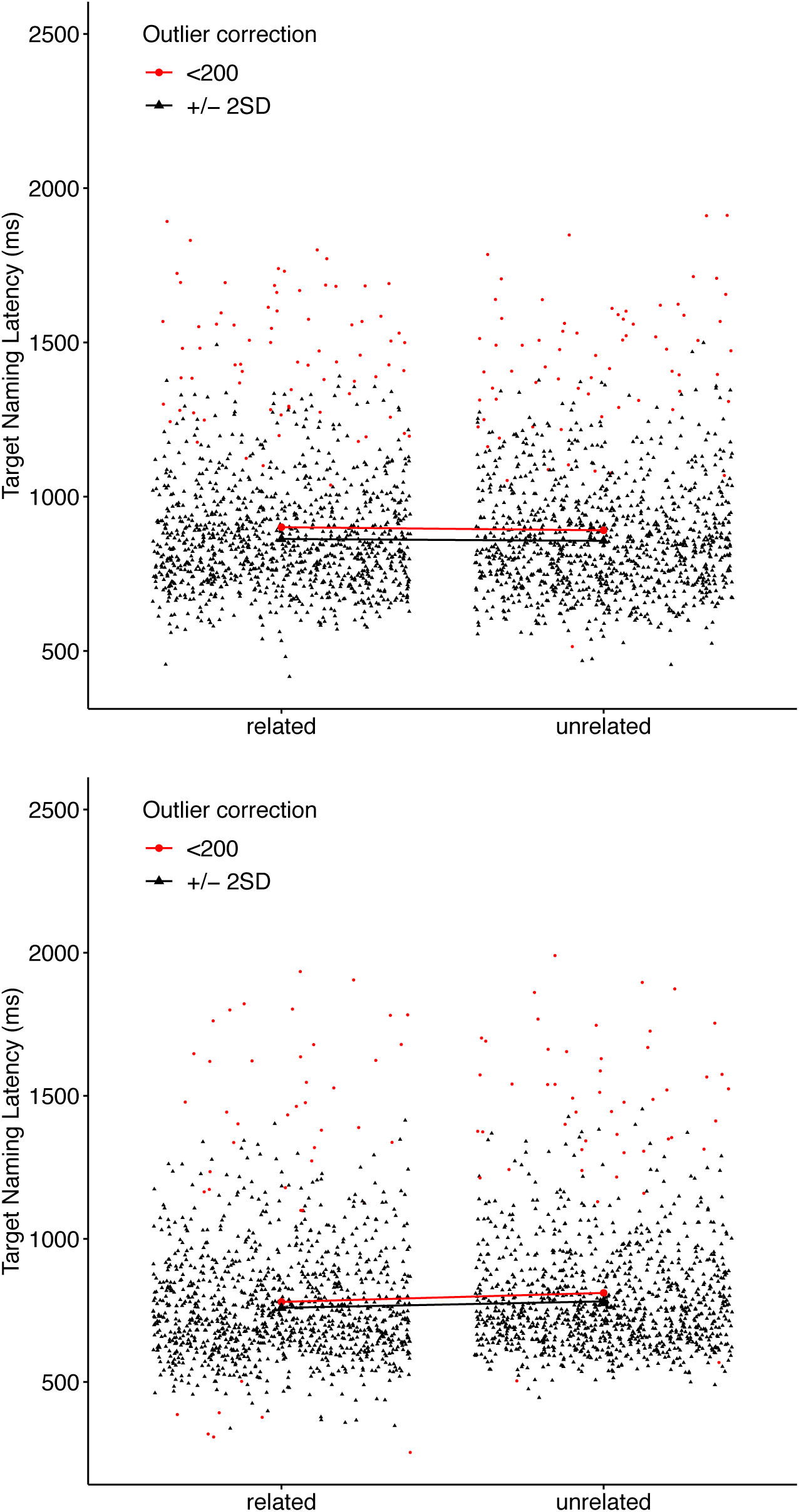
Experiment 3’s raw target naming latencies for each subject and item in the semantically related and unrelated condition for SOA-100 (top panel) and SOA-650 (bottom panel). Items marked as red circles would be excluded based on the pre- registered outlier correction (excluding trials +/-2SD of an individual’s mean). The implemented outlier correction (excluding trials <200ms) is less invasive.

## Appendix C

**Table C.1:** Preregistered model output for Experiment 1, testing an interaction between SOA and semantic relatedness. Fixed-effect estimates, standard error, 95% confidence intervals, t- values, and p-values based on Satterthwaites approximations. Model equation: log(target naming latency) ∼ SOA*semantic relatedness + (SOA| subject) + (semantic relatedness| target).

| *Predictors* | **Log-Transformed Target Naming Latencies** |  |  |  |  |
| --- | --- | --- | --- | --- | --- |
|  | *Estimates* | *SE* | *95% CI* | *t-value* | *p-value* |
| (Intercept) | 6.65 | 0.03 | 6.60 – 6.71 | 229.94 | <0.001 |
| SOA | -0.08 | 0.02 | -0.13 – -0.04 | -3.44 | 0.002 |
| Semantic Relatedness | 0.01 | 0.01 | -0.01 – 0.02 | 0.82 | 0.419 |
| SOA: Semantic Relatedness | 0.02 | 0.01 | -0.01 – 0.04 | 1.48 | 0.139 |
| **Random Effects** |  |  |  |  |  |
| Residual | 0.04 |  |  |  |  |
| Subject (intercept) | 0.02 |  |  |  |  |
| Subject (SOA) | 0.02 |  |  |  |  |
| Target (intercept) | 0.01 |  |  |  |  |
| Target (Semantic Relatedness) | 0.00 |  |  |  |  |
| Deviance | -1225.925 |  |  |  |  |
| log-Likelihood | 600.181 |  |  |  |  |

\* $p < 0.05$ \*\* $p < 0.01$ \*\*\* $p < 0.001$

**Table C.2:** Preregistered model output for Experiment 2, testing an interaction between SOA and semantic relatedness. Fixed-effect estimates, standard error, 95% confidence intervals, t- values, and p-values based on Satterthwaites approximations.. Model equation: log (target naming latency) ∼ SOA*semantic relatedness + (SOA| subject) + (semantic relatedness| target).

| *Predictors* | **Log-Transformed Target Naming Latencies** |  |  |  |  |
| --- | --- | --- | --- | --- | --- |
|  | *Estimates* | *SE* | *95% CI* | *t-value* | *p-value* |
| (Intercept) | 6.64 | 0.02 | 6.60 – 6.69 | 291.89 | <0.001 |
| SOA | -0.04 | 0.03 | -0.09 – 0.02 | -1.35 | 0.186 |
| Semantic Relatedness | -0.02 | 0.01 | -0.03 – 0.00 | -1.69 | 0.100 |
| SOA: Semantic Relatedness | 0.02 | 0.01 | -0.00 – 0.05 | 1.66 | 0.096 |
| **Random Effects** |  |  |  |  |  |
| Residual | 0.05 |  |  |  |  |
| Subject (intercept) | 0.01 |  |  |  |  |
| Subject (SOA) | 0.02 |  |  |  |  |
| Target (intercept) | 0.01 |  |  |  |  |
| Target (Semantic Relatedness) | 0.00 |  |  |  |  |
| Deviance | -224.737 |  |  |  |  |
| log-Likelihood | 99.671 |  |  |  |  |
| *$p < 0.05$ **$p < 0.01$ ***$p < 0.001$ | | | | | |

**Table C.3:** Preregistered model output for Experiment 3, testing an interaction between SOA and semantic relatedness. Fixed-effect estimates, standard error, 95% confidence intervals, t- values, and p-values based on Satterthwaites approximations. Model equation: log(target naming latency) ∼ SOA*semantic relatedness + (semantic relatedness*SOA| subject) + (semantic relatedness*SOA| target).

| **Log-Transformed Target Naming Latencies** |  |  |  |  |  |
| --- | --- | --- | --- | --- | --- |
| *Predictors* | *Estimates* | *SE* | *95% CI* | *t-value* | *p-value* |
| (Intercept) | 6.72 | 0.02 | 6.67 – 6.76 | 292.00 | <0.001 |
| SOA | -0.12 | 0.02 | -0.16 – -0.07 | -5.07 | <0.001 |
| Semantic Relatedness | 0.02 | 0.01 | -0.00 – 0.03 | 1.79 | 0.080 |
| SOA: Semantic Relatedness | 0.06 | 0.02 | 0.02 – 0.09 | 3.10 | 0.003 |
| **Random Effects** |  |  |  |  |  |
| Residual | 0.03 |  |  |  |  |
| Subject (intercept) | 0.01 |  |  |  |  |
| Subject (Semantic Relatedness) | 0.00 |  |  |  |  |
| Subject (SOA) | 0.02 |  |  |  |  |
| Subject (Semantic Relatedness: SOA) | 0.00 |  |  |  |  |
| Target (intercept) | 0.01 |  |  |  |  |
| Target (Semantic Relatedness) | 0.00 |  |  |  |  |
| Target (SOA) | 0.00 |  |  |  |  |
| Target (Semantic Relatedness: SOA) | 0.00 |  |  |  |  |
| Deviance | -2002.338 |  |  |  |  |
| log-Likelihood | 988.410 |  |  |  |  |

\* $p < 0.05$ \*\* $p < 0.01$ \*\*\* $p < 0.001$

## Appendix D

**Figure D.1.**
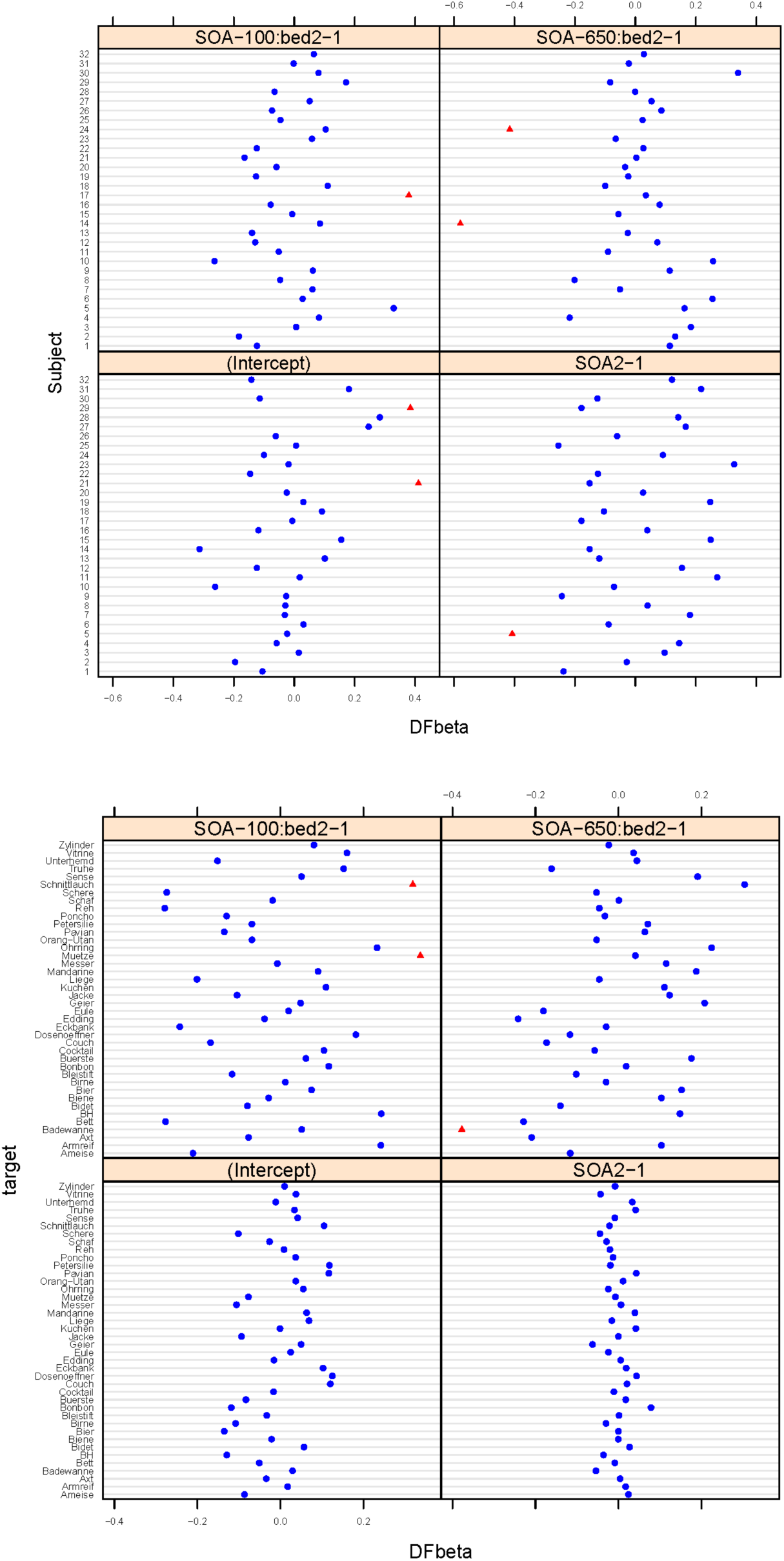
Dfbetas for participants (top panel) and target items (bottom panel) of Experiment 1. Red triangles mark overly influential participants or targets.

**Figure D.2.**
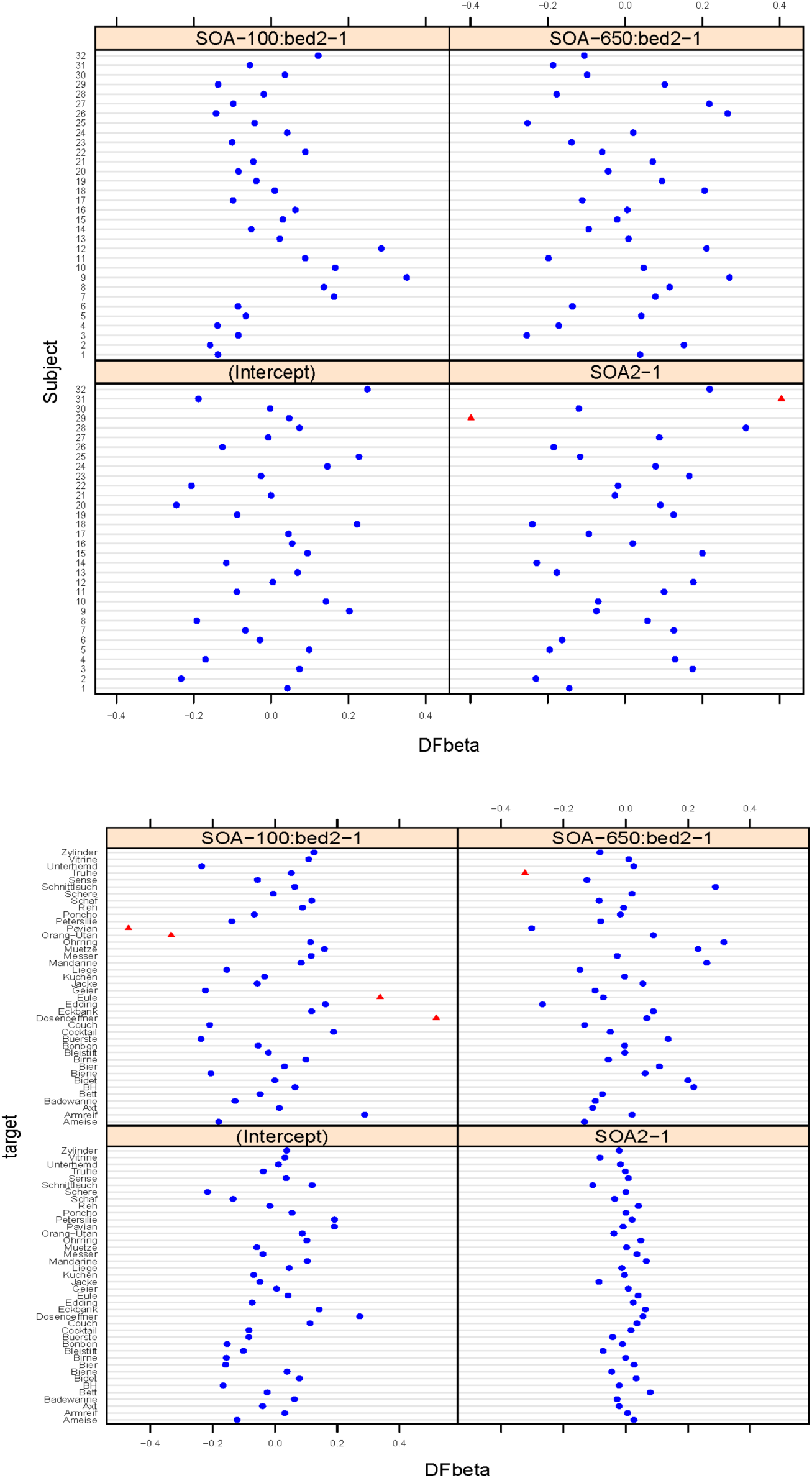
Dfbetas for participants (top panel) and target items (bottom panel) of Experiment 2. Red triangles mark overly influential participants or targets.

**Figure D.3.**
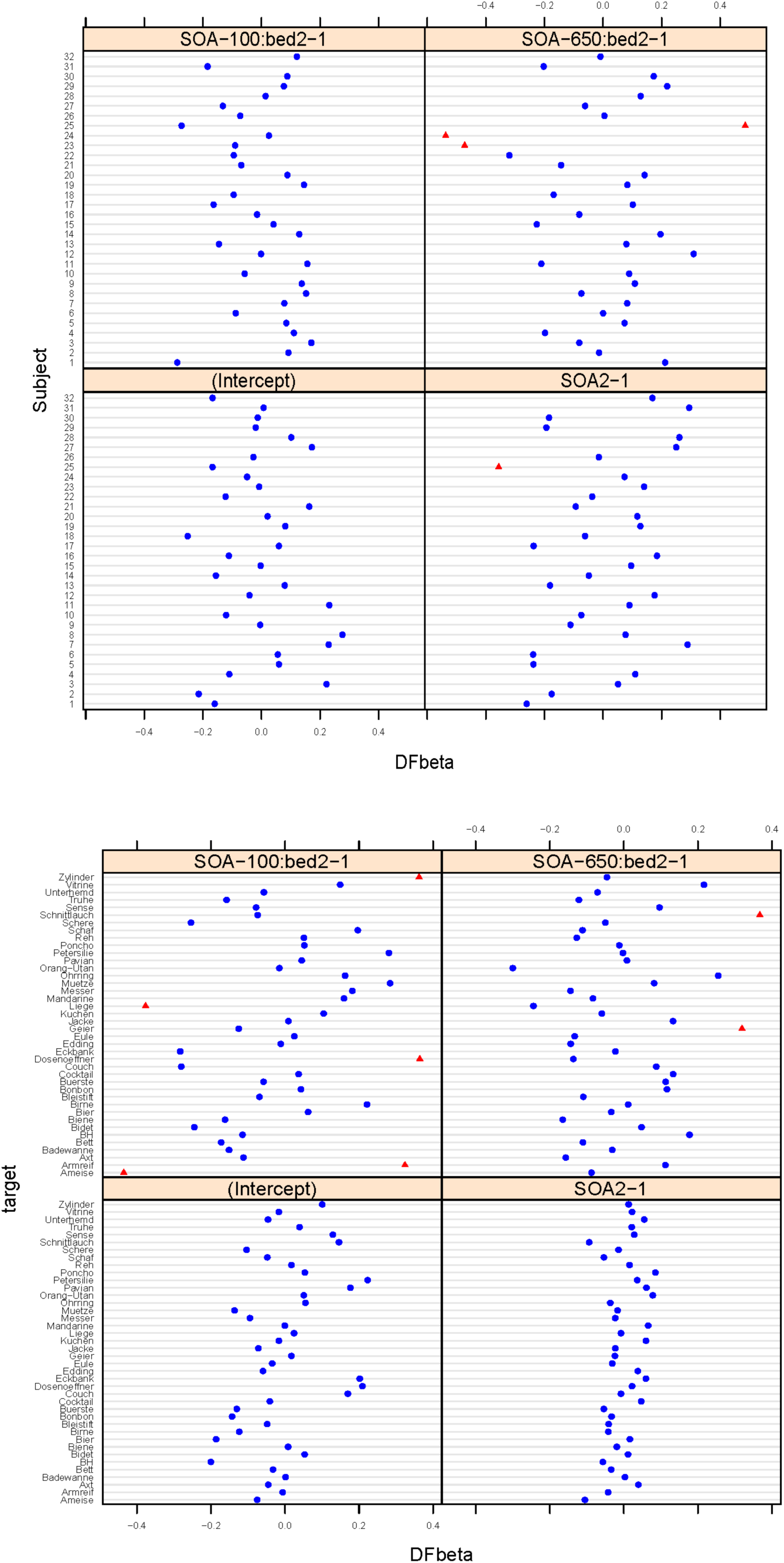
Dfbetas for participants (top panel) and target items (bottom panel) of Experiment 3. Red triangles mark overly influential participants or targets.

## Appendix F

**Table F.1:** Model output for LMM pooling data from Experiment 1 & 2 and entering word duration as predictor. Fixed-effect estimates, standard error, 95% confidence intervals, t- values, and p-values based on Satterthwaites approximations. Model equation: log (target naming latency) ∼ Experiment / ((SOA / semantic relatedness) + duration) + (SOA| subject) + (semantic relatedness| target)

| Log-Transformed Target Naming Latencies |  |  |  |  |  |
| --- | --- | --- | --- | --- | --- |
| *Predictors* | *Estimates* | *SE* | *95% CI* | *t-value* | *p-value* |
| (Intercept) | 6.65 *** | 0.02 | 6.61 – 6.69 | 330.53 | <0.001 |
| Exp2-1 | -0.01 | 0.03 | -0.07 – 0.05 | -0.32 | 0.753 |
| Exp1:SOA | -0.09 *** | 0.03 | -0.14 – -0.04 | -3.42 | 0.001 |
| Exp2:SOA | -0.04 | 0.03 | -0.09 – 0.01 | -1.54 | 0.128 |
| Exp1:Duration | 0.00 *** | 0.00 | 0.00 – 0.00 | 4.06 | <0.001 |
| Exp2:Duration | 0.00 *** | 0.00 | 0.00 – 0.00 | 6.25 | <0.001 |
| Exp1:SOA-100:Semantic Relatedness | -0.00 | 0.01 | -0.02 – 0.02 | -0.18 | 0.857 |
| Exp2:SOA-100:Semantic Relatedness | -0.03 * | 0.01 | -0.05 – -0.01 | -2.54 | 0.011 |
| Exp1:SOA-650:Semantic Relatedness | 0.02 | 0.01 | -0.01 – 0.04 | 1.45 | 0.147 |
| Exp2:SOA-650:Semantic Relatedness | -0.01 | 0.01 | -0.03 – 0.02 | -0.49 | 0.624 |
| **Random Effects** |  |  |  |  |  |
| Residual | 0.05 |  |  |  |  |
| Subject (intercept) | 0.01 |  |  |  |  |
| Subject (SOA) | 0.02 |  |  |  |  |
| Target (intercept) | 0.01 |  |  |  |  |
| Target (Semantic Relatedness) | 0.00 |  |  |  |  |
| Deviance | -1490.827 |  |  |  |  |
| log-Likelihood | 700.372 |  |  |  |  |

\* $p < 0.05$ \*\* $p < 0.01$ \*\*\* $p < 0.001$

## Appendix E

**Figure E.1.**
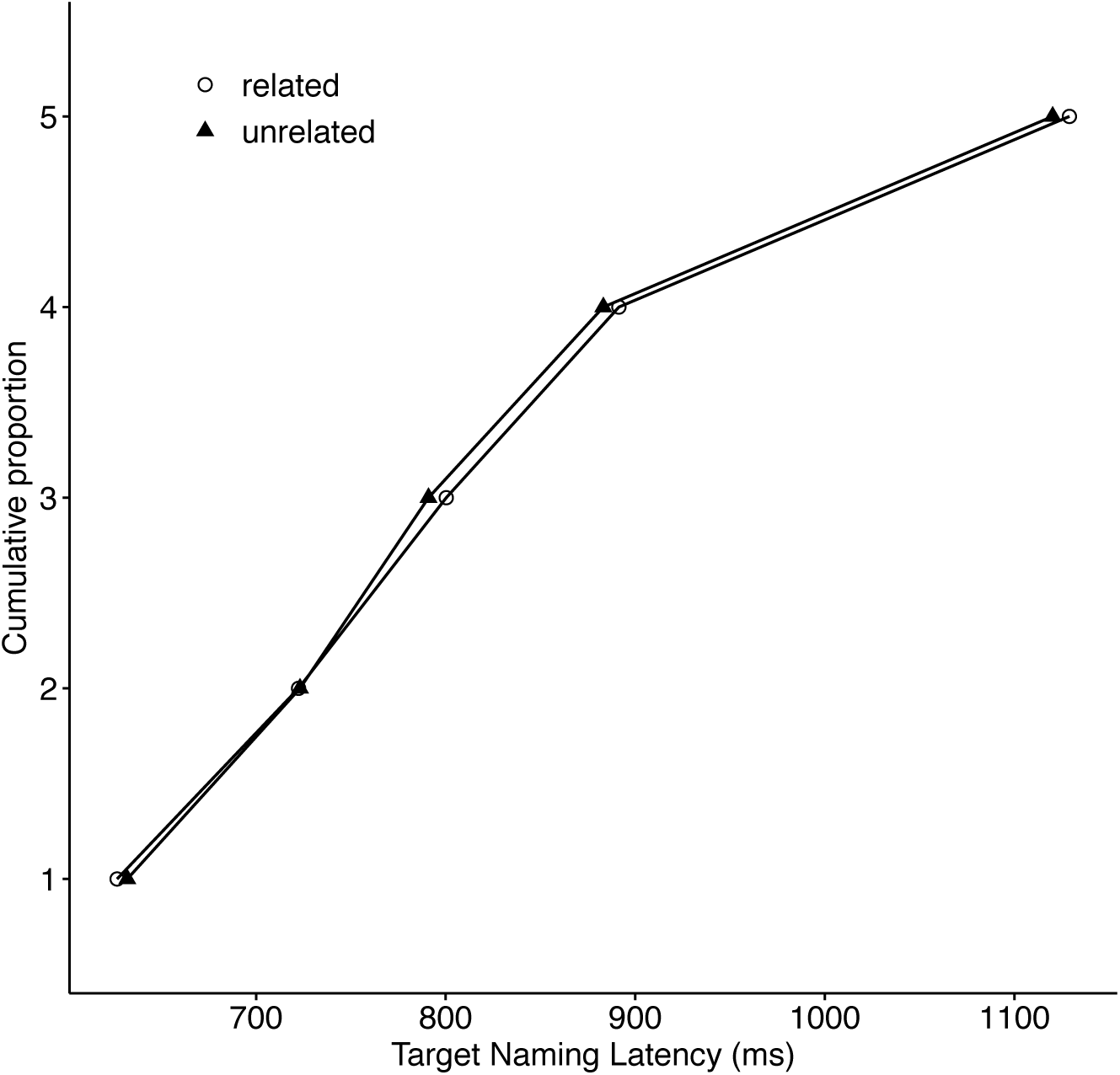
Vincentized distribution plots for Experiment 1.

**Figure E.2.**
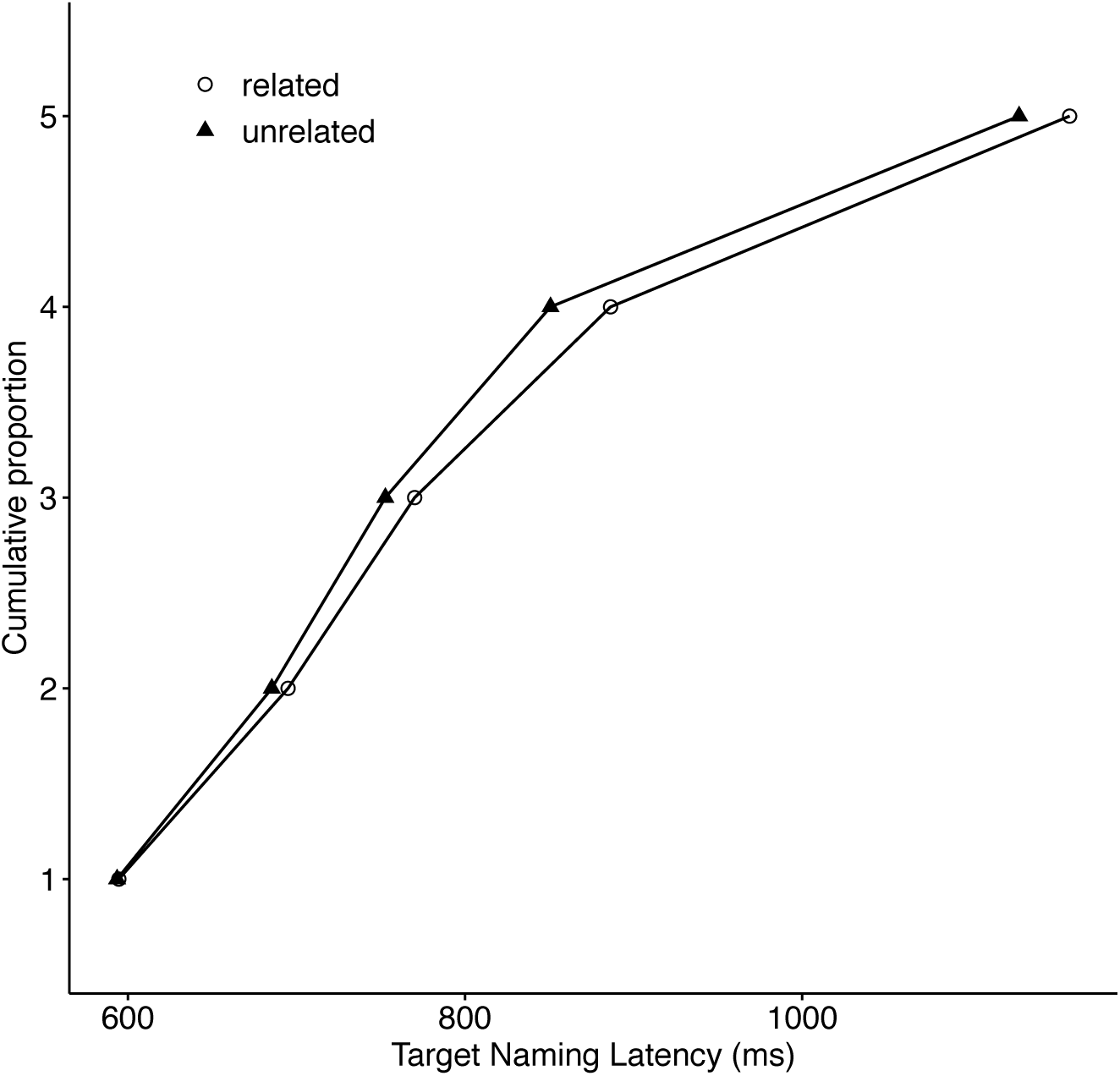
Vincentized distribution plots for Experiment 2.

**Figure E.3.**
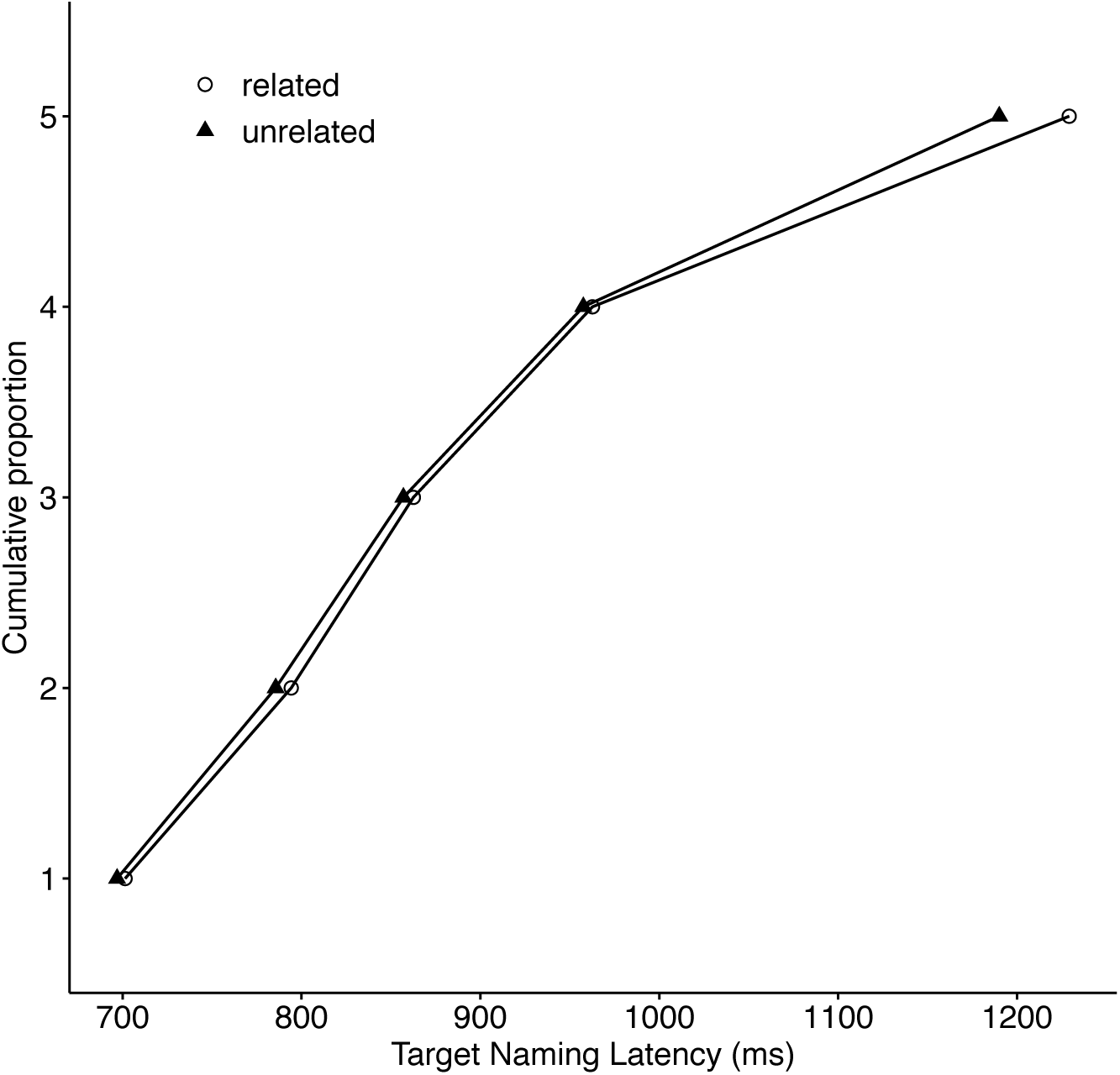
Vincentized distribution plots for Experiment 3.

## Footnotes

1 An analyses of the target naming latencies prior to excluding and replacing these participants yielded identical effects.

2 We pre-registered an outlier correction excluding naming latencies above two standard deviations of an individual’s overall mean. We now believe this may cut off differences between experimental conditions and the less invasive correction we report here is more truthful to data. We illustrate which trials would have been excluded following the preregistration in Appendix B. The decision to modify the outlier correction was made prior to analyzing our data. When implementing the pre-registered correction results stay the same with the pre- registered correction, with the exception of the interference effect at SOA-100 in Experiment 2, which reaches only marginally significant at p = .07. To address the possibility of individual subjects or items overly influencing our models we performed additional control analyses reported in the results.

## References

Abdel Rahman, R., & Melinger, A. (2007). When Bees Hamper the Production of Honey: Lexical Interference From Associates in Speech Production. Journal of Experimental Psychology: Learning, Memory and Cognition, 33(3), 604–614. doi:10.1037/0278-7393.33.3.604

Abdel Rahman, R., & Melinger, A. (2009). Semantic context effects in language production: A swinging lexical network proposal and a review. Language and Cognitive Processes, 24(5), 713–734. https://doi.org/10.1080/01690960802597250

Abdel Rahman, R. & Melinger, A. (2010). The dynamic microstructure of speech production: Semantic interference built on the fly. Journal of Experimental Psychology: Learning, Memory, and Cognition, 37(1), 149–161. doi: 10.1037/a0021208

Abdel Rahman, R., & Melinger, A. (2019). Semantic processing during language production: An update of the swinging lexical network. Language, Cognition and Neuroscience, 34(9), 1176–1192. https://doi.org/10.1080/23273798.2019.1599970

Alario, F. X., Segui, J., & Ferrand, L. (2000). Semantic and associative priming in picture naming. The Quarterly Journal of Experimental Psychology. A, Human Experimental Psychology, 53(3), 741–764. https://doi.org/10.1080/713755907

Allen, M., Poggiali, D., Whitaker, K., Marshall, T. R., & Kievit, R. A. (2019). Raincloud plots: A multi-platform tool for robust data visualization. Wellcome Open Research, 4. https://doi.org/10.12688/wellcomeopenres.15191.1

Baayen, R. H., Davidson, D. J., & Bates, D. M. (2008). Mixed-effects modeling with crossed random effects for subjects and items. Journal of Memory and Language, 59(4), 390–412. https://doi.org/10.1016/j.jml.2007.12.005

Bates, D., Kliegl, R., Vasishth, S., & Baayen, H. (2018). Parsimonious Mixed Models. ArXiv:1506.04967 [Stat]. http://arxiv.org/abs/1506.04967

Bates, D., Maechler, M., Bolker, B., & Walker, S. (2015). Fitting linear mixed-effects models using lme4. Journal of Statistical Software, 67(1), 1–48.

Baus, C., Sebanz, N., Fuente, V. de la, Branzi, F. M., Martin, C., & Costa, A. (2014). On predicting others’ words: Electrophysiological evidence of prediction in speech production. Cognition, 133(2), 395–407. https://doi.org/10.1016/j.cognition.2014.07.006

Belsley, D., Kuh, E., & Welsch, R. (1980). Regression Diagnostics. Identifying Influential Data and Sources of Collinearity. Wiley.

Bloem, I., van den Boogaard, S., & La Heij, W. (2004). Semantic facilitation and semantic interference in language production: Further evidence for the conceptual selection model of lexical access. Journal of Memory and Language, 51(2), 307– 323. https://doi.org/10.1016/j.jml.2004.05.001

Brehm, L., Taschenberger, L., & Meyer, A. (2019). Mental representations of partner task cause interference in picture naming. Acta Psychologica, 199, 102888. https://doi.org/10.1016/j.actpsy.2019.102888

Bürki, A., Elbuy, S., Madec, S., & Vasishth, S. (2020). What did we learn from forty years of research on semantic interference? A Bayesian meta-analysis. Journal of Memory and Language, 114, 104125. https://doi.org/10.1016/j.jml.2020.104125.

Clark, H. H. (1996). Using Language. Cambridge University Press.

Damian, M. F., & Martin, R. C. (1999). Semantic and phonological codes interact in single word production. Journal of Experimental Psychology. Learning, Memory, and Cognition, 25(2), 345–361. https://doi.org/10.1037//0278-7393.25.2.345

Du Bois, J. W., & Englebretson, R. (2004). SBC031 Tastes Very Special. Santa Barbara corpus of spoken American English, Part 3. Philadelphia: Linguistic Data Consortium. ISBN 1-58563-308-9.

Finkbeiner, M., & Caramazza, A. (2006). Now You See it, Now you Don’t: On Turning Semantic Interference Into Facilitation in a Stroop-Like Task. Cortex, 42(6), 790– 796. https://doi.org/10.1016/S0010-9452(08)70419-2

Forstmann, B. U., Jahfari, S., Scholte, H. S., Wolfensteller, U., van den Wildenberg, W. P. M., & Ridderinkhof, K. R. (2008a). Function and structure of the right inferior frontal cortex predict individual differences in response inhibition: A model-based approach. The Journal of Neuroscience, 28, 9790–9796. http://dx.doi.org/10.1523/JNEUROSCI.1465-08.2008

Gambi, C., Van de Cavey, J., & Pickering, M. J. (2015). Interference in joint picture naming. Journal of Experimental Psychology. Learning, Memory, and Cognition, 41(1), 1– 21. https://doi.org/10.1037/a0037438

Glaser, W. R., & Düngelhoff, F.-J. (1984). The time course of picture-word interference. Journal of Experimental Psychology: Human Perception and Performance, 10(5), 640–654. https://doi.org/10.1037/0096-1523.10.5.640

Green, MacLeod, & Nakagawa. (2016). SIMR: An R package for power analysis of generalized linear mixed models by simulation. Methods in Ecology and Evolution, 7(4), 493–498. https://doi.org/10.1111/2041-210X.12504

Hantsch, A., Jescheniak, J. D., & Schriefers, H. (2009). Distractor modality can turn semantic interference into semantic facilitation in the picture–word interference task: Implications for theories of lexical access in speech production. Journal of Experimental Psychology: Learning, Memory, and Cognition, 35(6), 1443–1453. https://doi.org/10.1037/a0017020

Hoedemaker, R. S., Ernst, J., Meyer, A. S., & Belke, E. (2017). Language production in a shared task: Cumulative Semantic Interference from self- and other-produced context words. Acta Psychologica, 172, 55–63. https://doi.org/10.1016/j.actpsy.2016.11.007

Hoedemaker, R. S., & Meyer, A. S. (2019). Planning and coordination of utterances in a joint naming task. Journal of Experimental Psychology. Learning, Memory, and Cognition, 45(4), 732–752. https://doi.org/10.1037/xlm0000603

Jescheniak, J. D., Kurtz, F., Schriefers, H., Günther, J., Klaus, J., & Mädebach, A. (2017). Words we do not say—Context effects on the phonological activation of lexical alternatives in speech production. Journal of Experimental Psychology: Human Perception and Performance, 43, 1194–1206.

Kessler, B., Treiman, R., & Mullennix, J. (2002). Phonetic Biases in Voice Key Response Time Measurements. Journal of Memory and Language, 47(1), 145–171. https://doi.org/10.1006/jmla.2001.2835

Kleinman, D., Runnqvist, E., & Ferreira, V. S. (2015). Single-word predictions of upcoming language during comprehension: Evidence from the cumulative semantic interference task. Cognitive Psychology, 79, 68–101. https://doi.org/10.1016/j.cogpsych.2015.04.001

Kuhlen, A. K., & Abdel Rahman, R. (2020, August 18). Social Picture-Word Interference. Retrieved from osf.io/3b9sr

Kuhlen, A. K., & Abdel Rahman, R. (2019, May 22). Social Picture-Word Interference. Pre-registration under AsPredicted.org, retrieved from http://aspredicted.org/blind.php?x=je2v8w

Kuhlen, A. K., & Abdel Rahman, R. (2017). Having a task partner affects lexical retrieval: Spoken word production in shared task settings. Cognition, 166, 94–106. https://doi.org/10.1016/j.cognition.2017.05.024

Kuznetsova, A., Brockhoff, P. B., & Christensen, R. H. B. (2017). lmerTest Package: Tests in Linear Mixed Effects Models. Journal of Statistical Software, 82(13). https://doi.org/10.18637/jss.v082.i13

Levelt, W. J., Roelofs, A., & Meyer, A. S. (1999). A theory of lexical access in speech production. The Behavioral and Brain Sciences, 22(1), 1–38

Levinson, S. C. (1983). Pragmatics. Cambridge University Press.

Lin, H. P., Kuhlen, A. K., & Abdel Rahman, R. (2021). Ad-hoc thematic relations form through communication: effects on lexical-semantic processing during language production. Language, Cognition and Neuroscience. DOI: 10.1080/23273798.2021.1900580

Lorenz, A., Regel, S., Zwitserlood, P., & Rahman, R. A. (2018). Age-related effects in compound production: Intact lexical representations but more effortful encoding. Acta Psychologica, 191, 289–309. https://doi.org/10.1016/j.actpsy.2018.09.001

Mädebach, A., Kurtz, F., Schriefers, H., & Jescheniak, J. D. (2020) Pragmatic constraints do not prevent the co-activation of alternative names: evidence from sequential naming tasks with one and two speakers. Language, Cognition and Neuroscience, DOI: 10.1080/23273798.2020.1727539

Mahon, B. Z., Costa, A., Peterson, R., Vargas, K. A., & Caramazza, A. (2007). Lexical selection is not by competition: A reinterpretation of semantic interference and facilitation effects in the picture-word interference paradigm*. Journal of Experimental Psychology: Learning*, Memory, and Cognition, 33(3), 503–535. https://doi.org/10.1037/0278-7393.33.3.503

Nieuwenhuis, R., te Grotenhuis, M., & Pelzer, B. (2012). Influence.ME: Tools for Detecting Influential Data in Mixed Effects Models. R Journal, 4(2), 38–47.

Pallier, C. (2002). Shuffle: A program to randomize lists with optional sequential constraints. Retrieved from http://www.pallier.org/lectures/shuffle/, Accessed date: May 15, 2020.

Piai, V., Roelofs, A., & Schriefers, H. (2011). Semantic interference in immediate and delayed naming and reading: Attention and task decisions. Journal of Memory and Language, 64, 404–423.

Proctor, R. W., Miles, J. D., & Baroni, G. (2011). Reaction time distribution analysis of spatial correspondence effects. Psychonomic Bulletin & Review, 18 (2), 242–266. doi:10.3758/s13423-011-0053-5

Ratcliff, R. (1979). Group reaction time distributions and an analysis of distribution statistics. Psychological Bulletin, 86(3), 446–461. https://doi.org/10.1037/0033-2909.86.3.446

Ridderinkhof, K. R. (2002). Activation and suppression in conflict tasks: Empirical clarification through distributional analyses. In W. Prinz & B. Hommel (Eds.), Attention and performance XIX: Common mechanisms in perception and action (pp. 494–519). New York: Oxford University Press.

Roelofs, A. (2008). Dynamics of the attentional control of word retrieval: Analyses of response time distributions. Journal of Experimental Psychology: General, 137(2), 303–323. https://doi.org/10.1037/0096-3445.137.2.303

Roelofs, A. (2018). A unified computational account of cumulative semantic, semantic blocking, and semantic distractor effects in picture naming. Cognition, 172, 59– 72. https://doi.org/10.1016/j.cognition.2017.12.007

Roelofs, A., & Piai, V. (2017). Distributional analysis of semantic interference in picture naming. The Quarterly Journal of Experimental Psychology 70 (4), 782–792. http://dx.doi.org/10.1080/17470218.2016.1165264

Rose, S. B., & Abdel Rahman, R. (2016). Cumulative semantic interference for associative relations in language production. Cognition, 152, 20–31. doi:http://dx.doi.org/10.1016/j.cognition.2016.03.013

Rose, S. B., Aristei, S., Melinger, A., & Abdel Rahman, R. (2019). The closer they are, the more they interfere: Semantic similarity of word distractors increases competition in language production. Journal of Experimental Psychology. Learning, Memory, and Cognition, 45(4), 753–763. https://doi.org/10.1037/xlm0000592

Sacks, H., Schegloff, E. A., & Jefferson, G. (1974). A Simplest Systematics for the Organization of Turn-Taking for Conversation. Language, 50(4), 696. https://doi.org/10.2307/412243

Scaltritti, M., Navarrete, E., & Peressotti, F. (2015). Distributional analyses in the picture-word interference paradigm: Exploring the semantic interference and distractor frequency effects. The Quarterly Journal of Experimental Psychology, 68, 1348– 1369.

Sellaro, R., Treccani, B., & Cubelli, R. (2018). When task sharing reduces interference: evidence for division-of labour in Stroop-like tasks. Psychological Research, 84(2), 327–342. doi: 10.1007/s00426-018-1044-1

Schriefers, H., Meyer, A. S., & Levelt, W. J. M. (1990). Exploring the time course of lexical access in language production: Picture-word interference studies. Journal of Memory and Language, 29(1), 86–102. https://doi.org/10.1016/0749-596X(90)90011-N

Shao, Z., & Rommers, J. (2019). How a question context aids word production: Evidence from the picture-word interference paradigm. Quarterly Journal of Experimental Psychology (2006), 73(2), 165–173. https://doi.org/10.1177/1747021819882911

Shao, Z., Meyer, A. S., & Roelofs, A. (2013). Selective and nonselective inhibition of competitors in picture naming. Memory & Cognition, 41(8), 1200–1211. https://doi.org/10.3758/s13421-013-0332-7

Shao, Z., Roelofs, A., Martin, R. C., & Meyer, A. S. (2015). Selective inhibition and naming performance in semantic blocking, picture-word interference, and color–word Stroop tasks. Journal of Experimental Psychology: Learning, Memory, and Cognition, 41(6), 1806–1820. https://doi.org/10.1037/a0039363

Sharma, D., Booth, R., Brown, R., & Huguet, P. (2010). Exploring the temporal dynamics of social facilitation in the Stroop task. Psychonomic Bulletin & Review, 17(1), 52– 58. https://doi.org/10.3758/PBR.17.1.52

Stivers, T., Enfield, N. J., Brown, P., Englert, C., Hayashi, M., Heinemann, T., Hoymann, G., Rossano, F., De Ruiter, J. P., Yoon, K. E., & Levinson, S. C. (2009) Universals and cultural variation in turn-taking in conversation. Proceedings of the National Academy of Sciences of the United States of America 106(26), 10587–92.

Tufft, M. R. A. & Richardson, D. (2020). Social offloading: just working together is enough to remove semantic interference. In S. Denison, M. Mack, Y. Xu, & B. Armstrong (Eds.), Proceedings of the 42nd Annual Conference of the Cognitive Science Society. Austin, TX: Cognitive Science Society Proceedings of the Cognitive Society.

van den Wildenberg, W. P. M., Wylie, S. A., Forstmann, B. U., Burle, B., Hasbroucq, T., & Ridderinkhof, K. R. (2010). To head or to heed? Beyond the surface of selective action inhibition: A review. Frontiers in Human Neuroscience, 4, 222. http://dx.doi.org/10.3389/fnhum.2010.00222

